# The cohesin loader NIPBL interacts with pre-ribosomal RNA and treacle to regulate ribosomal RNA synthesis

**DOI:** 10.1101/658492

**Authors:** Xiangduo Kong, Yen-Yun Chen, Jianhuang Lin, Ebony Flowers, Eric Van Nostrand, Steven M. Blue, Jonathan Chau, Christopher I-Hsing Ma, Isaiah Mohr, Ryan Thai, Chengguo Yao, Alexander R. Ball, Richard Chien, Shimako Kawauchi, Rosaysela Santos, Anne L. Calof, Arthur D. Lander, Yongsheng Shi, Mitsuru Okuwaki, Gene W. Yeo, Kyoko Yokomori

**Affiliations:** Department of Biological Chemistry, School of Medicine, University of California, Irvine, CA 92697, U.S.A.; PhD Program in Human Biology, School of Integrative and Global Majors and Faculty of Medicine, University of Tsukuba, Japan; California State University Long Beach, CA 90840, USA; Department of Cellular & Molecular Medicine and Institute for Genomic Medicine, University of California, San Diego, La Jolla, CA 92093, USA; Department of Microbiology & Molecular Genetics, School of Medicine, University of California, Irvine, CA 92697, U.S.A.; Department of Anatomy & Neurobiology, School of Medicine, University of California, Irvine, CA 92697, U.S.A.; Department of Developmental & Cell Biology, School of Biological Sciences, University of California, Irvine, CA 92697, U.S.A.; UT Southwestern Medical Center 5323 Harry Hines Blvd, NA8.124 Dallas, TX 75390; Department of Pathology, Anatomy, and Laboratory Medicine, West Virginia University, School of Medicine, Morgantown, WV 26506-9128

## Abstract

NIPBL is an essential loader of cohesin to mediate sister chromatid cohesion and chromatin loop organization. NIPBL mutations cause Cornelia de Lange Syndrome. How NIPBL’s genomic localization is specified is not fully understood. We found that NIPBL localizes to the nucleolus in an RNA-dependent manner and binds directly to ribosomal RNA (rRNA). We identified two RNA binding domains in NIPBL in vitro, both of which are required for efficient rRNA binding in vivo. NIPBL binds to ribosomal DNA (rDNA) in an RNA-stimulated manner, recruits PAF1 and promotes pre-rRNA transcription. Stress that inhibits rRNA synthesis displaces NIPBL from the nucleolus and rDNA. Interestingly, treacle, mutated in Treacher Collins syndrome, tightly binds to and recruits NIPBL to the nucleolus, nucleolar organizer regions, and the stress-induced nucleolar cap. The results reveal that a subpopulation of NIPBL is recruited to the nucleolus through its interaction with RNA and treacle and regulates pre-rRNA transcription.

## Introduction

NIPBL (or Scc2) is an evolutionarily conserved, essential protein that is required for chromatin loading of cohesin (Chien et al., 2011a; Dorsett, 2011; Horsfield et al., 2012). Cohesin is the essential multiprotein complex that plays a pivotal role in mediating chromatin interactions between sister chromatids for mitosis and DNA repair as well as chromosome domain organization and enhancer-promoter interactions affecting gene expression (Dekker and Mirny, 2016; Merkenschlager and Nora, 2016; Yokomori and Shirahige, 2017). Mutations in the *NIPBL* gene on chromosome 5p13 are linked to more than 55% of Cornelia de Lange Syndrome (CdLS) (OMIM 122470) cases, and NIPBL haploinsufficiency is associated with severe cases of CdLS (Krantz et al., 2004; Tonkin et al., 2004). CdLS is an autosomal dominant multisystem developmental disorder characterized by facial dysmorphism, hirsutism, upper limb abnormalities, cognitive retardation, and growth abnormalities (Boyle et al., 2015; DeScipio et al., 2005; Liu and Krantz, 2009). NIPBL haploinsufficiency does not overtly affect sister chromatid cohesion, but reduces cohesin binding-mediated chromatin interactions and alters gene expression, which appear to directly contribute to the developmental phenotype of the disease (Chien et al., 2011b; Muto et al., 2011; Newkirk et al., 2017; Remeseiro et al., 2013). Evidence also suggests that NIPBL may have cohesin-independent functions in gene regulation (Zuin et al., 2014), consistent with the notion that NIPBL haploinsufficiency often exhibits more severe and somewhat distinct phenotypes from CdLS and CdLS-like disorders caused by cohesin subunit mutations (OMIM300590 (CdLS1 (*SMC1A*)), 610759 (CdLS3 (*SMC3*)), 614701 (CdLS4 (*RAD21*))) (Deardorff et al., 2007; Deardorff et al., 2012; Mannini et al., 2013; Musio et al., 2006). Several factors have been shown to mediate NIPBL loading to specific genomic regions, including the pre-replication complex (for the replication origins), Mediator (for the active promoter/enhancer regions), HP1 (for H3 lysine 9 methylated heterochromatin), and a neuronal transcription factor/integrator complex (for RNA polymerase II (pol II) paused sites in neuronal cells) (Gillespie and Hirano, 2004; Kagey et al., 2010; Lechner et al., 2005; Takahashi et al., 2004; van den Berg et al., 2017; Zeng et al., 2009a). Considering the variety of NIPBL/cohesin binding sites, either constitutive or cell type-specific, it would be expected that there may be other mechanisms and interacting partners that dictate NIPBL (and cohesin) binding specificity. These, however, are not fully elucidated.

The nucleolus is a sub-nuclear compartment where ribosomal RNA (rRNA) synthesis and ribosome assembly take place, which is tightly linked to cell growth and proliferation (Boulon et al., 2010b; Grummt, 2013). In addition, the nucleolus plays a pivotal role in viral replication, signal recognition particle assembly, and modification of small nuclear RNPs (Pederson, 2011). Thus, the nucleolus is considered a central hub for RNA metabolism. The nucleolus also serves as a critical sensor for stress, and alters its structure and protein composition in response. Since ribosome biosynthesis is a highly energy-consuming process, virtually any types of stress result in alteration of the nucleolar structure and inhibition of rRNA synthesis to limit cell cycle and growth and preserve energy required to maintain cellular homeostasis (Boulon et al., 2010b; Grummt, 2013). Mutations of genes that impair rRNA transcription or ribosome biogenesis have been found in a group of developmental disorders termed “ribosomopathies” (Hannan et al., 2012; Teng et al., 2013). These include Treacher Collins Syndrome, Bloom’s and Werner Syndromes, Cockayne Syndrome, and Shwachman–Diamond Syndrome. Although mutations of these genes all impair aspects of ribosome biogenesis, they are linked to tissue-specific distinct disease phenotypes (Yelick and Trainor, 2015).

The ribosomal DNA (rDNA) repeats were one of the first cohesin binding sites identified in yeast (Laloraya et al., 2000), and we previously demonstrate that both cohesin and NIPBL bind to the rDNA regions in human cells (Zeng et al., 2009a). Smc1 mutation caused decreased ribosomal biogenesis and protein translation (Bose et al., 2012). Scc2 (yeast NIPBL homolog) was also found to affect rRNA transcription (Zakari et al., 2015). However, NIPBL’s role in ribosomal biogenesis in mammalian cells was unclear, and how NIPBL/Scc2 is recruited to the rDNA region was not understood. Here we report that NIPBL localizes specifically to the fibrillar center (FC) of the nucleolus where active RNA pol I transcription takes place and stimulate pre-rRNA synthesis through the recruitment of the PAF1 complex during interphase. This localization is mediated through interaction with rRNA and the treacle protein, the product of the *TCOF1* gene mutated in Treacher Collins Syndrome (TCS). In response to stresses, NIPBL dissociates from rDNA and relocalizes to the nucleolar cap in a treacle-dependent manner, contributing to the timely suppression of rRNA synthesis. Our results demonstrate that NIPBL is an RNA binding protein, and provide the first example of the transcript-dependent recruitment of NIPBL to specific genomic regions to regulate the gene expression. Our results also reveal an interesting potential link between CdLS and TCS.

## Results

### NIPBL localizes to the Fibrillar Center of the nucleolus

The nucleolus has a “tripartite structure” consisting of the Fibrillar Center (FC), the Dense Fibrillar Component (DFC), and the Granular Component (GC), each with distinct functions (Fig. 1A) (Boulon et al., 2010a). The FC is the site for pre-rRNA transcription and is enriched in RNA polymerase I (Pol I) machinery. Pre-rRNA is processed in the DFC, which contains many RNA processing factors such as snoRNA and fibrillarin. The GC is the site where pre-ribosome assembly takes place, and contains ribosome assembly factors such as B23 (nucleophosmin). Consistent with cohesin binding to the rDNA region (Zeng et al., 2009a), biochemical fractionation analysis indicates that Nipbl and cohesin are present in both nuclear and nucleolar fractions (Fig. 1B). Using an affinity-purified polyclonal antibody directed against the C-terminus of human NIPBL, we observed clear localization of both mouse Nipbl and human NIPBL in the nucleolus (mouse embryonic fibroblasts (MEFs) in Fig. 1C and D; human 293T cells in Supplemental Fig. S1, respectively). Costaining with antibodies specific for B23 (a marker for the GC) and fibrillarin (a marker for the DFC) revealed that Nipbl tightly clusters to the inside of the DFC corresponding to the FC (Fig. 1C and D). Furthermore, nucleolar Nipbl staining signals were reduced significantly in a Nipbl heterozygous mutant MEFs (modeling NIPBL haploinsufficiency in CdLS (Kawauchi et al., 2009; Newkirk et al., 2017)) and Nipbl siRNA-depleted MEFs compared to the wild type and control siRNA-treated MEFs, respectively (Fig. 1E). These results confirmed the specificity of immunostaining and also indicate that Nipbl in the nucleolus is sensitive to partial protein depletion (haploinsufficiency).

**Figure 1.**
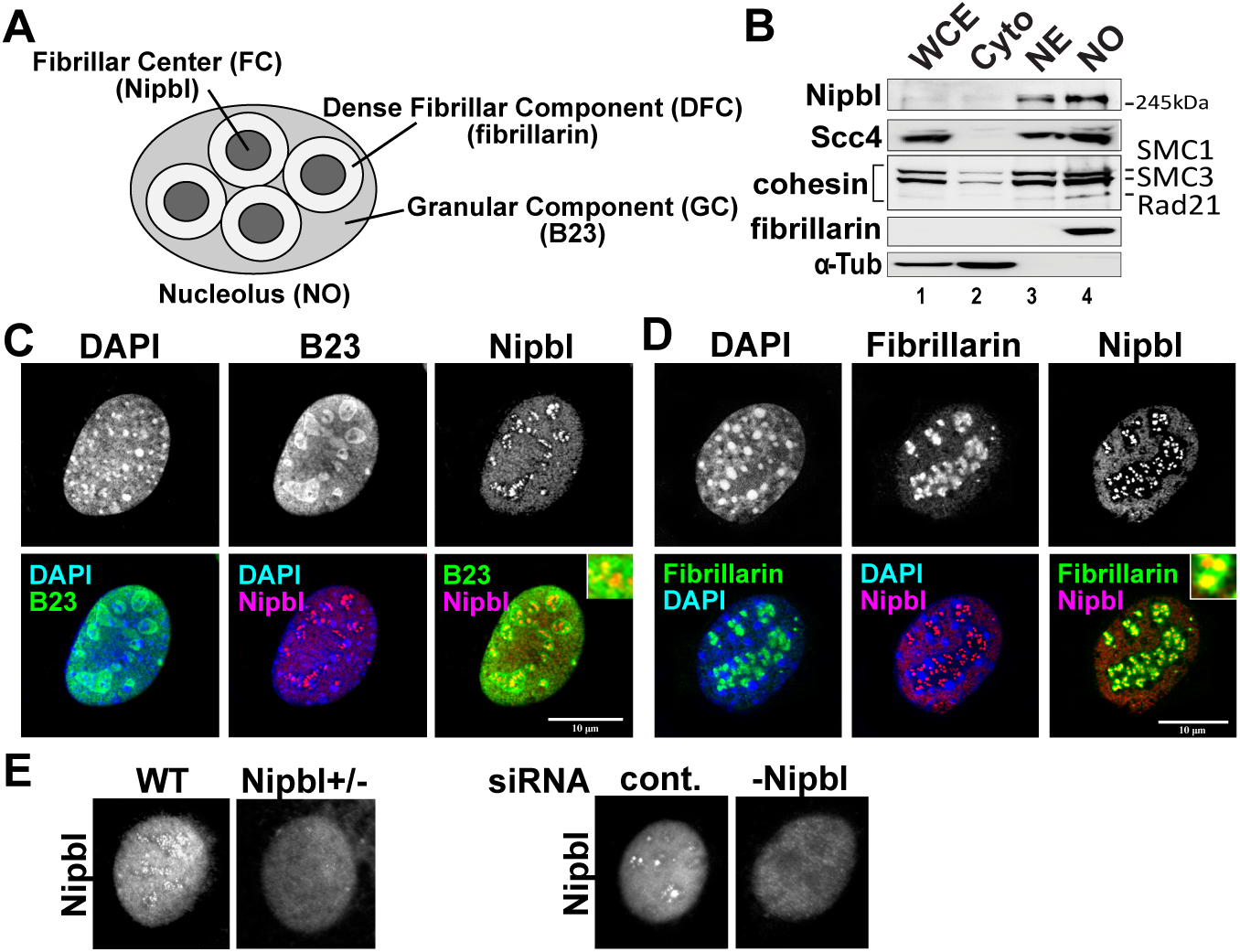
Nipbl forms foci in the FC region of the nucleolus. **A.** Schematic diagram of the tripartite structure in the nucleolus. **B.** Western blot analysis of whole cell (WCE), cytoplasmic (Cyto), nuclear (NE) and nucleolar (NO) fractions with antibodies specific for the indicated proteins. Nipbl and its binding partner Scc4 as well as cohesin are enriched in both NE and NO fractions. **C.** and D. Localization of Nipbl foci in the fibrillar center (FC) region in the nucleolus. Immunofluorescent staining of Nipbl, B23 (C) and fibrillarin (D) as indicated. Merged images including DAPI staining are shown underneath. Insets show higher magnification. Scale bar=10 μm. **E.** The effects of Nipbl reduction on the immunofluorescent signals of Nipbl foci in the nucleolus, indicating the specificity of the antibody. Nipbl signals in the nucleoli were reduced by partial reduction of Nipbl in heterozygous mutant (Nipbl+/-) (left), or in Nipbl siRNA-treated (-Nipbl) (right) MEFs. Three different wild type and Nipbl+/- MEF lines as well as cells transfected with control siRNA or two different Nipbl siRNAs were examined. Quantification of Nipbl transcripts is shown in Supplemental Table S1, demonstrating comparable reduction of Nipbl between heterozygous mutation and siRNA treatment.

### NIPBL localizes to the nucleolus in an RNA-dependent manner and binds to pre-ribosomal RNA

RNA is highly enriched and is an important constituent of nucleoli. Interestingly, RNase treatment dispersed Nipbl nucleolar foci while bulk Nipbl remained in the nucleoplasm (Fig. 2A). The results indicate that NIPBL localizes to the nucleolus in an RNA-dependent manner. This raises the possibility that Nipbl interacts with RNA. By in vivo RNA immunoprecipitation after UV crosslinking (CLIP), we detected robust binding of both 18S and 28S rRNAs by Nipbl (Fig. 2B). Nipbl depletion by two different siRNAs reduced the binding signals, confirming the specificity of the IP (Fig. 2B and Supplemental Fig. S2A). Rad21, a non-SMC subunit of cohesin, was previously shown to bind to estrogen receptor-transcribed enhancer RNA in human breast cancer cells (Li et al., 2013). Rad21 depletion failed to affect Nipbl binding to rRNA, suggesting that Nipbl binding to RNA is not mediated by cohesin (Supplemental Fig. S2B). Consistent with this, only a very weak rRNA binding signal was detected with Rad21 CLIP (Fig. 2B).

**Figure 2.**
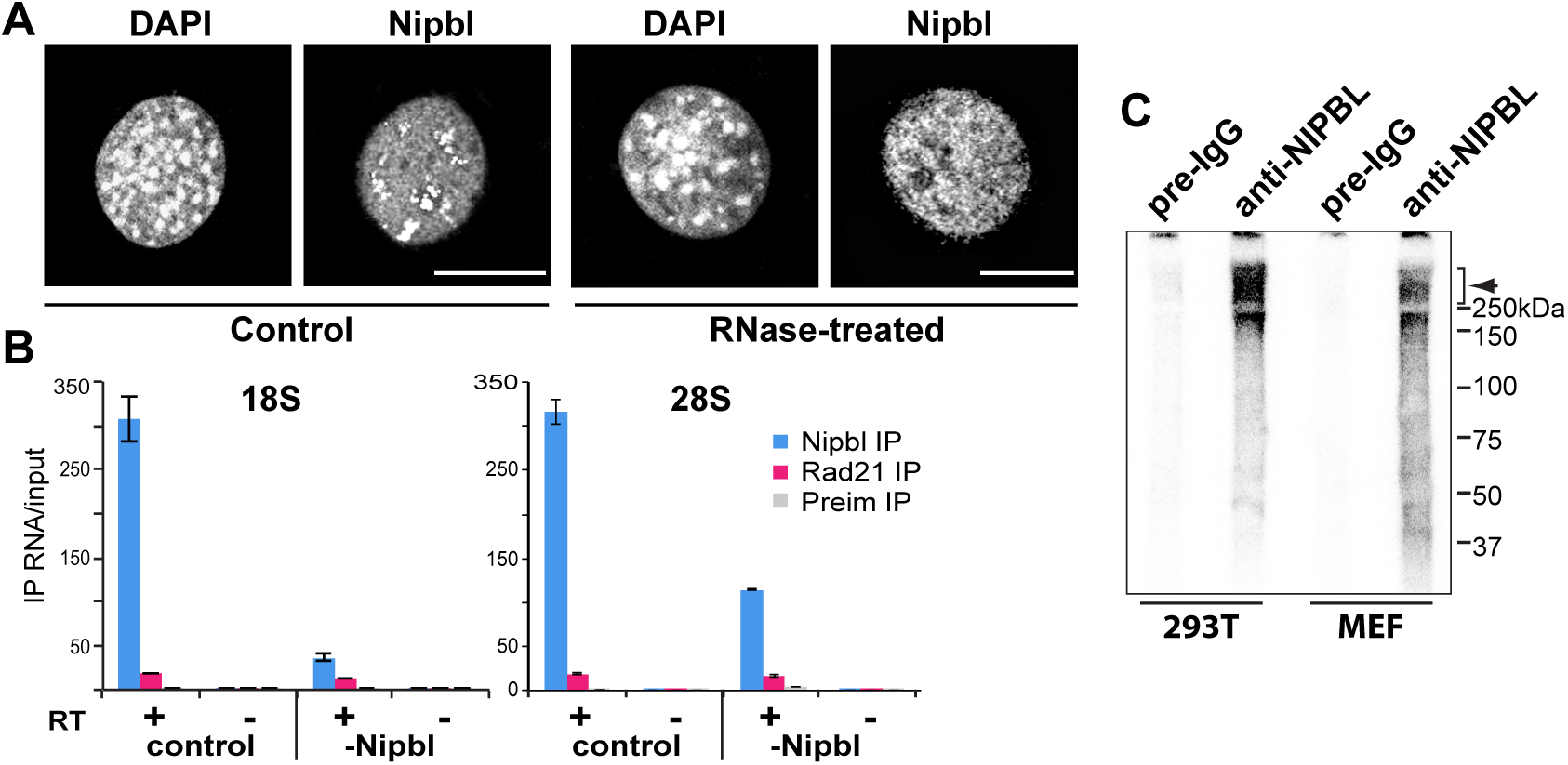
Nipbl localizes to the nucleolus in an RNA-dependent manner and binds to rRNA. **A.** RNase treatment disperses Nipbl from nucleoli. Immunofluorescent staining of control (left) and RNase-treated (right) MEFs with indicated antibodies. Scale bar=10 μm. **B.** Nipbl binds to rRNA. RNA crosslinking and immunoprecipitation (CLIP) followed by RT-qPCR using primers specific for 18S and 28S coding regions was performed either with (+) or without (-) reverse transcription (RT) in control or Nipbl siRNA-treated cells using antibodies specific for NIPBL or Rad21, or preimmune IgG. A schematic diagram of the rDNA locus and corresponding pre-rRNA transcript (45S), and the positions of PCR primers are shown underneath. Lack of PCR signals without RT indicates that the signals come from rRNA and not rDNA. Treatment with Nipbl siRNA reduced the binding signal. **C.** The iCLIP preparatory SDS-PAGE (CLIP/SDS-PAGE) using preimmune IgG (pre-IgG) or anti-NIPBL antibody using human 293T cells and MEFs. Crosslinked and immunoprecipitated RNAs using preimmune IgG or anti-NIPBL antibody were radio-labeled and subjected to SDS-PAGE. Radioactive signals (smear indicated by a bracket and arrowhead) above 250kDa are specific to Nipbl IP. Similar results were obtained using both human 293T cells and MEFs. Two different regions of 28S were recovered from RNA extracted from the bracketed region of the gel (Supplemental Fig. S2C).

Following CLIP, protein species that are directly bound by RNA can be resolved by SDS-PAGE following ^32^P end-labeling of crosslinked RNA (Chan et al., 2014). The radiolabeled protein species in NIPBL CLIP exhibited the size-range expected for the full-length and various isoforms and breakdown products of NIPBL/Nipbl proteins in both human and mouse cells (Zuin et al., 2014) (Fig. 2C). No other significant radiolabeling was observed corresponding, for example, to the NIPBL binding partner Mau2/Scc4 (∼66kDa) (Fig. 2C) (Watrin et al., 2006). Two separate sequences from 28S RNA were retrieved from RNA purified from the region above 250kDa in the gel (Supplemental Fig. S2C). Thus, the results indicate that NIPBL/Nipbl directly binds to rRNA in vivo.

To provide higher resolution mapping of NIPBL binding across the 45S ribosomal RNA precursor, we performed eCLIP to profile NIPBL RNA binding sites in K562 cells (Van Nostrand et al., 2016) (Fig. 3A and B). Analysis of eCLIP reads, including both unique genomic-mapped as well as rRNA (and other repetitive elements)-mapped reads, revealed an average 1.5-fold enrichment for NIPBL binding to rRNA, and in particular a 3-fold enrichment for 45S pre-rRNA (Fig. 3C). Mapping read density specifically across the 45S pre-rRNA, we observed substantial enrichment in NIPBL eCLIP at multiple loci compared to 126 other RNA binding proteins profiled by the ENCODE consortium (Fig. 3D) (Van Nostrand et al., 2016). Under our condition, we observed striking enrichment in the 5’ETS and regions flanking the 18S rRNA region (Fig. 3D), confirming that the enriched 45S signal is specific to NIPBL eCLIP. Consistent with the results above (Supplemental Fig. 2C), there is also specific enrichment of peaks in the 28S coding region (Fig. 3D). These results revealed statistically significant enrichment of pre-rRNA species in NIPBL eCLIP, and indicate that NIPBL/Nipbl specifically binds to pre-rRNA species in vivo.

**Figure 3.**
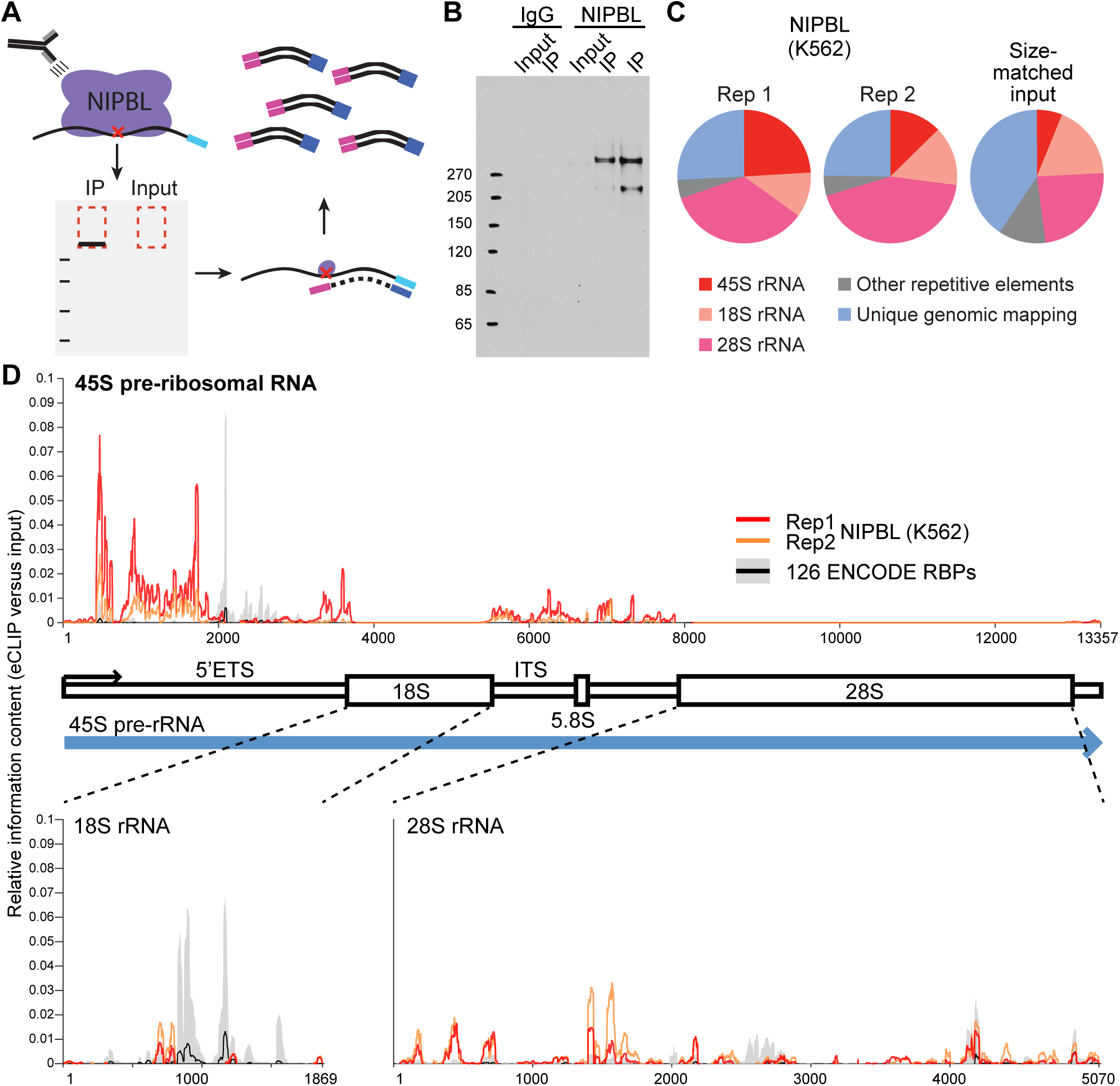
eCLIP indicates specific binding of NIPBL to pre-ribosomal RNA. **A.** Schematic of NIPBL eCLIP. After UV crosslinking, NIPBL is immunoprecipitated along with bound RNA and stringently washed. Following 3’ adapter ligation (blue), IP samples (along with a paired input) were subjected to PAGE electrophoresis and transferred to nitrocellulose membranes, and the region from NIPBL (estimated size 316 kDa) to 75 kDa above was extracted and treated with Proteinase K to isolate RNA. RNA was reverse transcribed into cDNA, a second adapter was ligated to the 3’ end, and samples were PCR amplified to obtain final libraries for sequencing. **B.** IP-western performed during eCLIP of NIPBL on 10% of IP sample (either using anti-NIPBL antibody or rabbit IgG isotype control) and 2% of total lysis as input. **C.** Pie charts indicate the fraction of total reads in NIPBL eCLIP that map to the indicated target categories. **D.** NIPBL enrichment at 45S pre-rRNA (top) and 18S and 28S rRNA (bottom). Red (replicate 1) and orange (replicate 2) lines indicate relative information in NIPBL IP versus input at each position along the indicated transcript, whereas the black line indicates the mean and grey box indicates standard deviation observed across 181 eCLIP experiments in K562 and HepG2 cells (profiling 126 total RBPs) performed by the ENCODE consortia.

### Characterization of NIPBL RNA binding domains

Sequence analysis using BindN (Wang and Brown, 2006) suggested potential RNA binding domains in NIPBL at the N- and C-termini (Fig. 4A, open boxes; Supplemental Fig. S3A). Thus, the RNA binding activity of these two domains was further examined in vitro. GST fused to N- and C-terminal fragments containing the putative RNA binding domains (Fig. 4A, NRBD and CRBD, respectively, indicated by black bars) were expressed in E. coli, purified on glutathione beads, and incubated with total RNA from 293T cells. Bound RNA species (without crosslinking) were analyzed by either RT-qPCR or visualized directly using two different gel systems (Fig. 4B). By RT-qPCR analysis, we found that both NRBD and CRBD bind to 18S and 28S as well as 5’ETS and ITS present only in unprocessed pre-rRNA species, consistent with the eCLIP results of the endogenous NIPBL (Figs. 3 and 4C; Supplemental Fig. S4). In contrast, GST fused to GFP or the C-terminal region of Rad21 exhibited no significant binding (Fig. 4C). Separating the CRBD into two putative RNA binding domains identified by BindN resulted in complete loss of the 18S and 28S RNA binding activity (with the reduced residual binding activity of C-CRBD to 5’ETS and ITS regions), suggesting that both regions are necessary for efficient rRNA binding (Supplemental Fig. S5).

**Figure 4.**
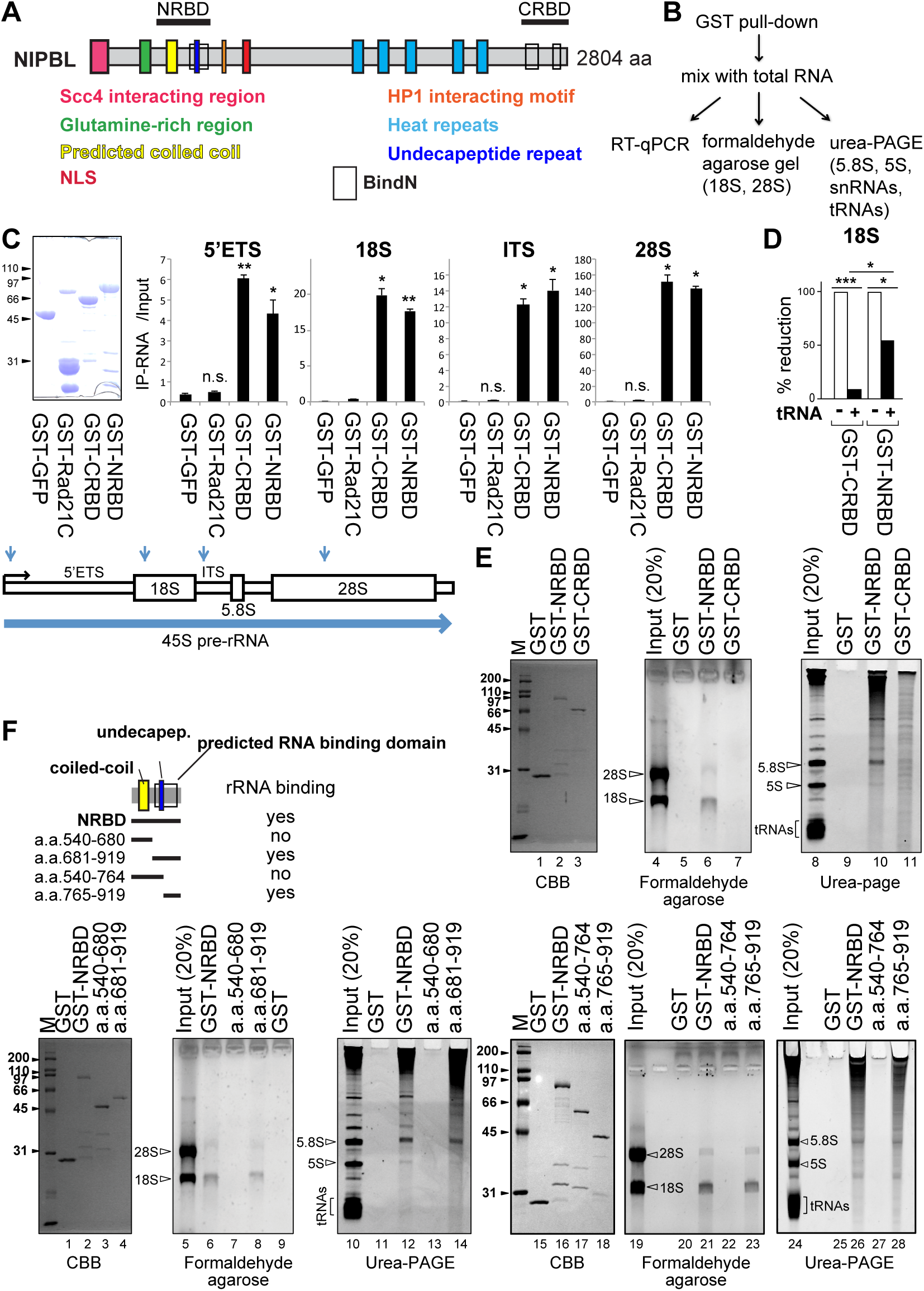
Identification of the RNA binding domains of NIPBL. **A.** A schematic diagram of NIPBL with known domains and the region containing putative RNA binding motif. In addition to the previously defined domains (colored boxes), putative RNA binding domains identified by BindN are indicated by open boxes. Two regions expressed in vitro (NRBD and CRBD) are indicated. **B.** Flow chart of in vitro RNA binding experiments. **C.** Left: Coomassie staining of recombinant purified GST fusion proteins (GST-GFP, GST- Rad21C, GST-CRBD and GST-NRBD). These proteins are used for in vitro RNA binding experiments on the right. Different regions (5’ETS, 18S, ITS and 28S) of rRNA were detected by PCR. Positions of PCR products in the rRNA are indicated underneath. GST-GFP and GST- Rad21C were used as negative controls. Error bars denote the standard deviation from the mean of three biological replicates and p-values were calculated by comparing to the GST-GFP negative control (* p<=0.05; ** p<=0.01; ns, not significant, p>0.05). **D.** RNA binding of GST-CRBD and -NRBD in the presence of excess tRNAs competed the binding of CRBD to 18S. Only partial reduction was seen with NRBD. * p<=0.05; *** p<=0.001. **E.** Similar experiments as in **(C)** except the bound RNA species were analyzed on formaldehyde agarose gel stained with EtBr (lanes 4-7) to analyze binding of 18S and 28S rRNA and Urea-PAGE stained with GelRed (lanes 8-11) for 5.8S, 5S and tRNA species. The left panel (lanes 1- 3): Coomassie staining (CBB) of GST, GST-NRBD and GST-CRBD used in the binding experiments. **F.** Similar experiments as in (**E**) with the NRBD deletion mutants: a.a. 540-680 containing coiled-coil region, a.a. 681-919, including the predicted RNA binding domain coinciding with the undecapeptide repeat region, a.a. 540-764, including coiled-coil and undecapeptide repeat regions but missing the C-terminal half of the predicted RNA binding domain, and a.a. 765-919, including the C-terminal region just downstream of the undecapeptide region. Lanes 1-4: Coomassie staining (CBB) of GST, GST- a.a. 540-680, and GST-a.a. 681-919. Lanes 5-9: formaldehyde agarose; lanes 10-14: urea-PAGE. Lanes 15-18: Coomassie staining (CBB) of GST, GST-NRBD, GST-a.a. 540-764, and GST-a.a. 765-919. Lanes 19-23: formaldehyde agarose; lanes 24-28: urea-PAGE. Schematic diagrams of deletion mutants are shown above.

Interestingly, addition of excess tRNA competitor almost completely inhibited rRNA binding of CRBD, while it only partially affected NRBD binding, suggesting that NRBD more specifically binds to rRNA (Fig. 4D). Consistent with this, direct analysis of bound RNA species (without PCR amplification) revealed that clear interaction of NRBD, but not CRBD, with 18S and 28S, with preferential enrichment of 18S (Fig. 4E). In urea-PAGE gel, NRBD appears to also bind 5.8S but not tRNAs, whereas modest and relatively non-specific RNA binding was detected with CRBD (Fig. 4E). NRBD contains a coiled-coil region and the putative RNA binding domain that overlaps with the undecapeptide repeats (Mannini et al., 2013) (Fig. 4F; Supplemental Fig. S3A). Deletion analysis revealed that the RNA binding activity resides in the C-terminal half whereas the coiled-coil region has no RNA binding activity (Fig. 4F, lanes 1-14). Further deletion analysis revealed that the robust RNA binding activity is in the charged domain (a.a. 765-919) (isoelectric point (pI)=10.3) just downstream of the undecapeptide repeats (a.a. 514-764) (pI=6.6) (Fig. 4F, lanes 15-28). This region has no known motif and is expected to contain intrinsically disordered regions (IDRs) (Supplemental Fig. S3B). Recent evidence suggests that not only classical structured domains but also IDRs with a large net charge function in RNA binding (Calabretta and Richard, 2015). Taken together, the results indicate that NIPBL contains at least two separate RNA binding domains with different specificities.

To substantiate the role of NRBD and CRBD in vivo, either the two fragments were expressed separately (499-1124 a.a. and 1844-2804 a.a., respectively) or the minimum domains were linked together (NRBD+CRBD) with a FLAG tag (Fig. 5A). With the comparable protein expression, NRBD+CRBD exhibited robust RNA binding activity compared to the two domains separately. Although much lower compared to the two domains combined, a slightly higher binding activity of the NRBD-containing domain was observed compared to the fragment containing CRBD only, consistent with the in vitro data (Figs. 4D and 5A). Conversely, deletion of NRBD and CRBD from the full-length NIPBL (ΔNRBD-ΔCRBD) significantly reduced the rRNA binding activity (Fig. 5B). Since residual RNA binding was still observed with ΔNRBD-Δ CRBD, possible contribution of additional domain(s) of NIPBL cannot be excluded. Nevertheless, our results indicate that both NRBD and CRBD are required for efficient rRNA binding of NIPBL in vivo. Interestingly, these domains appear to be conserved in human, mice, chicken and Xenopus, but not in drosophila, C. elegans or yeast (Fig. 5C).

**Figure 5.**
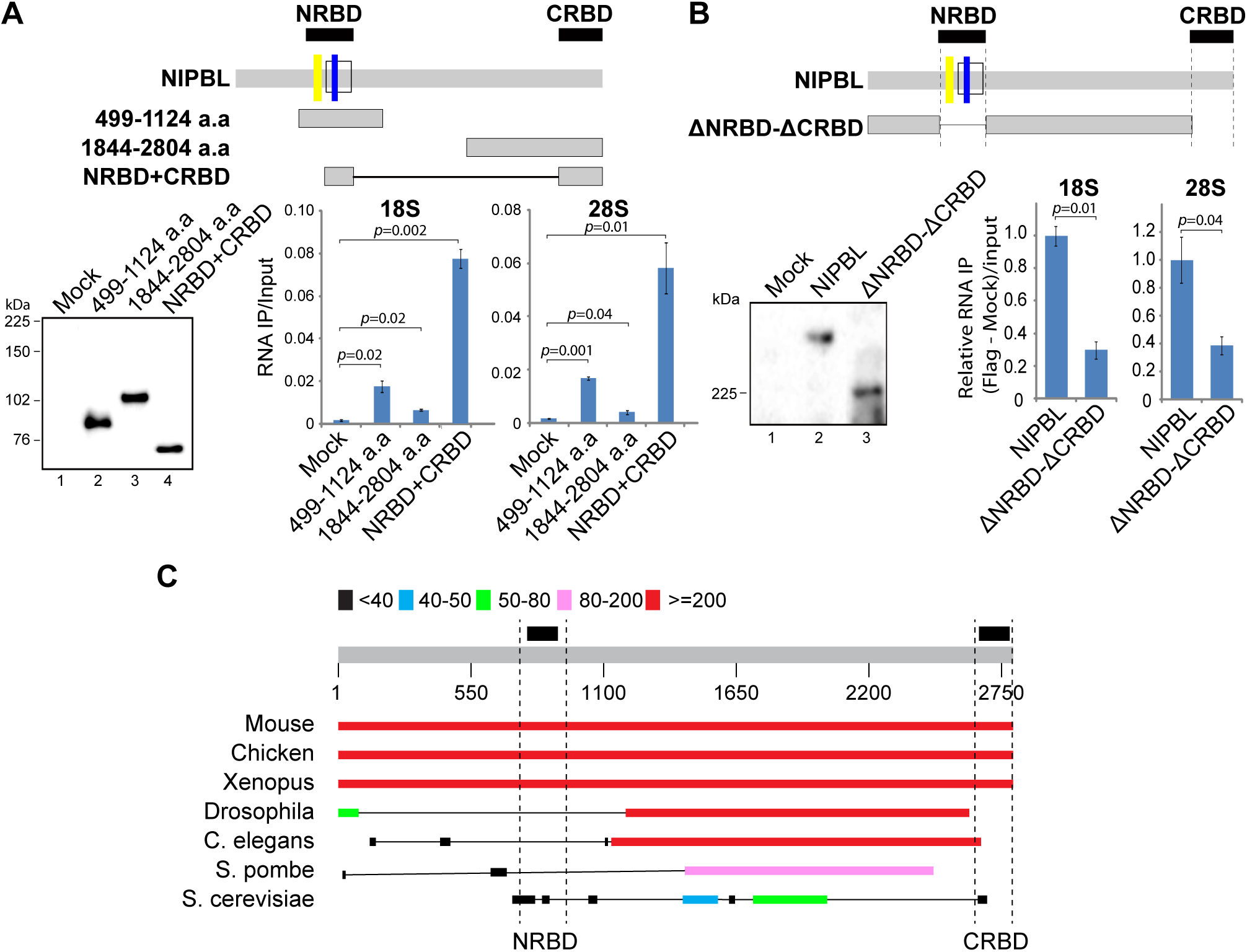
NRBD and CRBD are required for efficient NIPBL binding to rRNA in vivo. **A.** Top: Schematic diagrams of three NIPBL mutant constructs tested. Mutants are tagged with FLAG at the N-terminus and expressed in 293T cells. Bottom left: Western blot analysis of three mutant protein expression probed with anti-FLAG antibody. Bottom right: CLIP assay of mock- or mutant construct-transfected cells using anti-FLAG antibody. Precipiated RNA was analyzed by PCR using primers specific for 18S or 28S as in Figure 2B. Data is the mean ± SD of three biological replicates. **B.** Top: A schematic diagram of the NIPBL mutant lacking NRBD and CRBD (ΔNRBD-ΔCRBD). Bottom left: western analysis of mock, full-length NIPBL or mutant-transfected 293T cells. Bottom right: CLIP assay as in (A). Data is the mean ± SD of three biological replicates. **C.** NIPBL homolog protein alignment using protein BLAST (https://blast.ncbi.nlm.nih.gov/). The location of the NRBD and CRBD are indicated at the top (black boxes). Color coding for the alignment score is indicated.

### NIPBL binds to rDNA in an RNA-dependent manner

We next examined Nipbl binding to rDNA by ChIP-qPCR analysis in MEFs. Similar to our previous observation in human cells (Zeng et al., 2009a), we found that Nipbl binds preferentially to the rRNA coding region of DNA (18S and 28S), rather than the intergenic spacer regions (Fig. 6A). Although cohesin binds to the rDNA region in a NIPBL/Nipbl-dependent manner in mammalian cells (Zeng et al., 2009a), we observed that Nipbl and cohesin binding is discordant: Nipbl binding to the rDNA repeats is much more robust than at cohesin binding sites containing a CTCF motif in the RNA pol II gene regions (*Ebf1* and *Cebpβ*) whereas cohesin (and CTCF) binding is more significant at the latter (Fig. 6A). Since Nipbl clustering to the nucleoli requires RNA, we tested whether RNA is necessary for Nipbl association with rDNA. RNase treatment decreased Nipbl binding to the rDNA regions, but not to the unique cohesin binding sites (*Ebf1-In*, *Cebpβ* and *Cebpδ* promoter regions) (Fig. 6B). In contrast, B23’s association with rDNA was not affected by RNase treatment under this condition (Fig. 6B). These results indicate that the rRNA gene region is a major binding site for NIPBL, consistent with the localization of NIPBL in the NORs/FC region in the nucleolus, and the Nipbl binding to the rDNA region is uniquely dependent on RNA.

**Figure 6.**
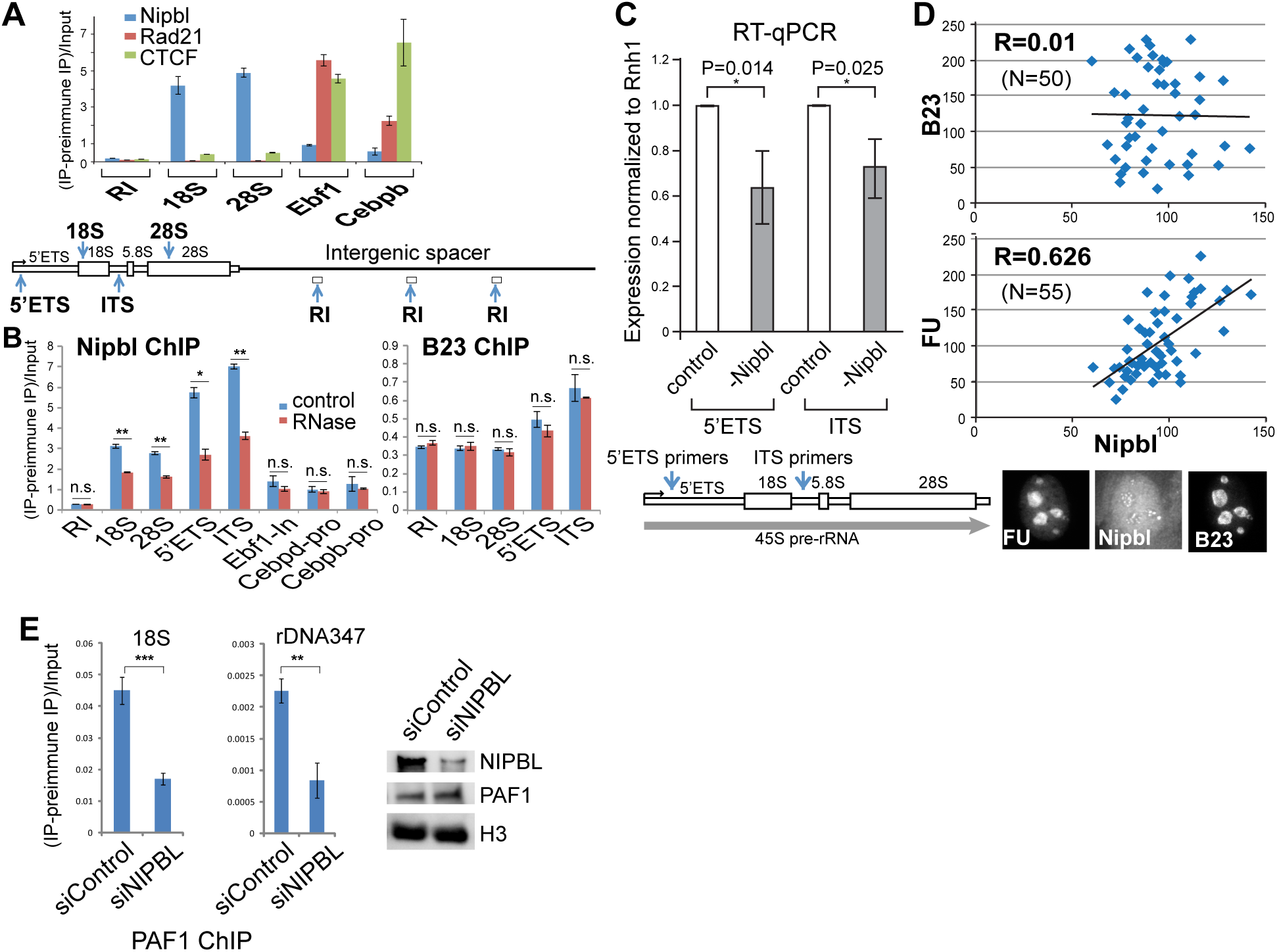
NIPBL binds to rDNA in an RNA-dependent manner and stimulates pre-rRNA synthesis and PAF1 recruitment. **A.** Disproportionate binding of Nipbl, cohesin, and CTCF at rDNA and unique cohesin binding sites. ChIP was performed using antibody specific for NIPBL, Rad21, and CTCF and the ChIP-DNA was analyzed by PCR using primers specific for the indicated regions. A schematic diagram of rDNA region with corresponding PCR products is shown. Cohesin/CTCF binding sites on Ebf1 and Cebpβ gene loci were examined for comparison. **B.** Similar ChIP-PCR analyses as in (A) using antibodies specific for NIPBL and B23 with and without RNase treatment. Nipbl binding to rRNA regions was specifically reduced by the treatment, but not to the unique cohesin binding sites in the pol II genes. B23 ChIP is not affected by RNase treatment. * p<=0.05; ** p<=0.01; ns, not significant, p>0.05. **C.** Analysis of the effect of Nipbl depletion on pre-rRNA expression. RT-qPCR analysis of pre-rRNA transcript in control- and Nipbl-siRNA-treated MEFs. Results were averaged from four independent experiments. PCR primers for the 5’ETS and ITS regions were used to detect pre-rRNA as shown below. **D.** Correlation between de novo rRNA synthesis and Nipbl localization in the nucleolus. Positive correlation between Nipbl foci and the nascent transcript signal (FU) detected in situ, but not with B23. Fluorescence intensity measurement of FU, Nipbl and B23 staining was performed in the same cells and their correlation was plotted. Top: Quantification of Nipbl nucleolar staining versus B23 nucleolar staining (N=50). Bottom: Quantification of Nipbl nucleolar staining versus FU labeling signals (N= 55). Fluorescent signals are normalized by the nucleolar areas. An example of a triple-stained cell nucleus is shown below. **E.** ChIP analysis of PAF1 binding to rDNA in control and NIPBL siRNA-treated cells. Western blot with indicated antibodies is shown to confirm that NIPBL depletion did not affect the PAF1 protein level. ** p<=0.01; *** p<=0.001.

### NIPBL recruits PAF1 and stimulates pre-rRNA synthesis

In yeast, loss of function mutation of Scc2 (NIPBL homolog) reduced pre-rRNA synthesis (Zakari et al., 2015). Consistent with this, we found that partial depletion of Nipbl in MEFs resulted in significant reduction of pre-rRNA transcripts (5’ETS and ITS) (Fig. 6C). Although CTCF was previously shown to bind to the rDNA promoter region and stimulate rRNA synthesis (Huang et al., 2013), we failed to observe any effect of CTCF depletion in the same assay (Supplemental Fig. S6). Nascent rRNA transcripts can be detected by incorporation of 5- fluorouridine (FU) in the nucleoli (Boisvert et al., 2000). The intensity of FU labeling correlated with Nipbl signal intensity (R=0.626) (Fig. 6D). This is in contrast to the lack of correlation between Nipbl and B23 signals (R=0.01). The results support that Nipbl localized in the nucleolus is involved in pre-rRNA synthesis.

The RNA polymerase-associated factor 1 complex (Paf1C) was shown to promote rRNA transcription elongation in yeast (Zhang et al., 2009), and Scc2 mutation compromised the recruitment of Paf1 to the rDNA region in S. cerevisiae (Zakari et al., 2015). We found that NIPBL depletion reduced PAF1 binding to rDNA region in human cells (Fig. 6E). Thus, the results suggest that RNA-dependent recruitment of NIPBL promotes further recruitment of PAF1 to stimulate pre-rRNA transcription.

### NIPBL localizes to the nucleolar cap in response to stress

The above results suggest that the rRNA transcript promotes Nipbl recruitment to the rDNA region, which in turn stimulates rRNA transcription. Conversely, we found that inhibition of rRNA transcription in response to stress displaced Nipbl from the nucleolus (Fig. 7). A low concentration of actinomycin D (ActD) inhibits Pol I transcription creating transcriptional stress, and relocalizes FC/ DFC components to nucleolar caps (Shav-Tal et al., 2005). When MEFs were treated with ActD, Nipbl nucleolar foci indeed became clustered to the “nucleolar cap” structure (Fig. 7A). Fibrillarin, a DFC protein, also relocalized to caps in response to ActD treatment; while B23, a protein in the GC region, was unaffected under this condition (Supplemental Fig. S7A). H_2_O_2_ and INK127 induce oxidative and nutrient stresses, respectively, which also induce Nipbl nucleolar cap formation (Fig. 7B). In human myoblast cells, H_2_O_2_ treatment significantly reduced NIPBL in the nucleolus (instead of forming a cap structure) (Supplemental Fig. S7B). Consistent with the displacement of NIPBL/Nipbl from the FC region, a significant decrease of NIPBL binding to the rDNA region was observed by ChIP analysis in H_2_O_2_-treated cells compared to control cells (Fig. 7C). This is in contrast to the NIPBL binding to other genomic regions, including heterochromatic repeats (D4Z4) and unique gene regions (*MYC-P2* and *FBXL16*), which was unaffected (Fig. 7C). These results indicate that NIPBL/Nipbl is part of the RNA Pol I silencing pathway central to the nucleolar stress response in mammalian cells.

**Figure 7.**
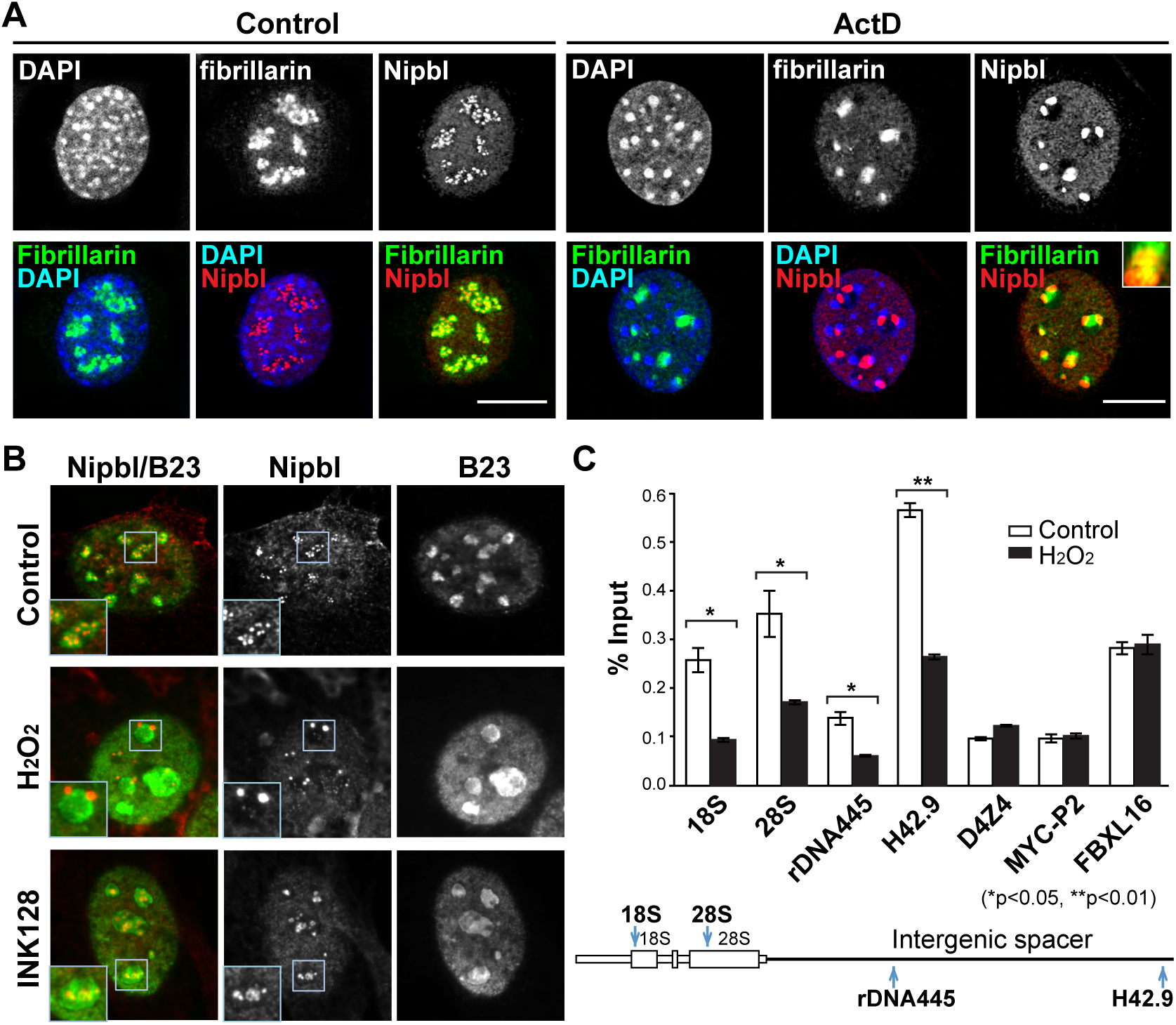
NIPBL relocalizes to the nucleolar cap structure in response to stress. **A.** The effect of Actinomycin D treatment on Nipbl localization in the nucleolus. MEFs were treated with Actinomycin D (Act D) and were stained with DAPI and antibodies specific for fibrillarin and NIPBL. **B.** The effects of H_2_O_2_ and INK128 treatment on the nucleolar localization of Nipbl. Nipbl moves to the nucleolar periphery (cap) by H_2_O_2_ treatment. With INK128, Nipbl clusters within the nucleolus. **C.** NIPBL binding to the rDNA region is specifically reduced by H_2_O_2_ treatment. NIPBL ChIP-qPCR analysis was performed in control and H2O2-treated human cells. The locations of PCR products in the rDNA region are indicated underneath. NIPBL binding to D4Z4 repeats, MYC-P2, and FBXL16, known cohesin/NIPBL binding sites, is not affected by H_2_O_2_ treatment. Data is the mean ± SD of three biological replicates, where statistical significance was calculated by comparing H_2_O_2_ treatment to control at each region using a Student’s two-tailed t-test. * p<=0.05; ** p<=0.01.

### A subpopulation of NIPBL tightly interacts with treacle

Tight localization of NIPBL/Nipbl in the FC region of the nucleolus and its effect on rRNA transcription prompted us to examine the NIPBL interaction with the additional factors involved in ribosomal RNA transcription. The Treacher Collins Syndrome *TCOF1* gene product treacle stimulates rRNA synthesis (Valdez et al., 2004). Treacle also binds to UBF, the pol I transcription activator (Valdez et al., 2004). Thus, we examined the NIPBL interaction with treacle and UBF. NIPBL closely colocalizes with treacle as well as UBF not only in interphase nucleoli but also during mitosis at nucleolar organizer regions (NORs), strongly suggesting that NIPBL is a primary component of the nucleoli (Fig. 8A; Supplemental Fig. S8). Antibody specific for NIPBL specifically co-precipitated both treacle and UBF (Fig. 8B, top). Interestingly, the NIPBL-treacle interaction, but not NIPBL-UBF, was resistant to a high salt wash, suggesting that NIPBL primarily interacts with treacle (Fig. 8B, top). No significant treacle signal was detected, however, in the cohesin subunit Rad21 coimmunoprecipitation, indicating that NIPBL, and not cohesin, is the primary interactor of treacle (Fig. 8B, bottom). RNase has no effect on the NIPBL-treacle interaction, and the ΔNRBD-ΔCRBD mutant still interacts with treacle, indicating that the NIPBL-treacle interaction is not mediated by RNA and the observed NIPBL RNA binding is not mediated by treacle (Fig. 8C and Supplemental Fig. S9A). In response to stress, treacle and NIPBL co-cluster to the nucleolar cap (Fig. 8A, bottom; Supplemental Fig. S9B). SiRNA depletion of treacle abolished NIPBL localization both to the FC region and to the nucleolar cap, indicating that the treacle interaction is critical for the NIPBL clustering in the nucleolus (Fig. 8A, middle and bottom; Supplemental Fig. S10). Taken together, the results indicate the unique combined mechanism of RNA- and treacle-dependent NIPBL loading to the nucleolus and rDNA loci to contribute to the regulation of ribosomal RNA biosynthesis.

**Figure 8.**
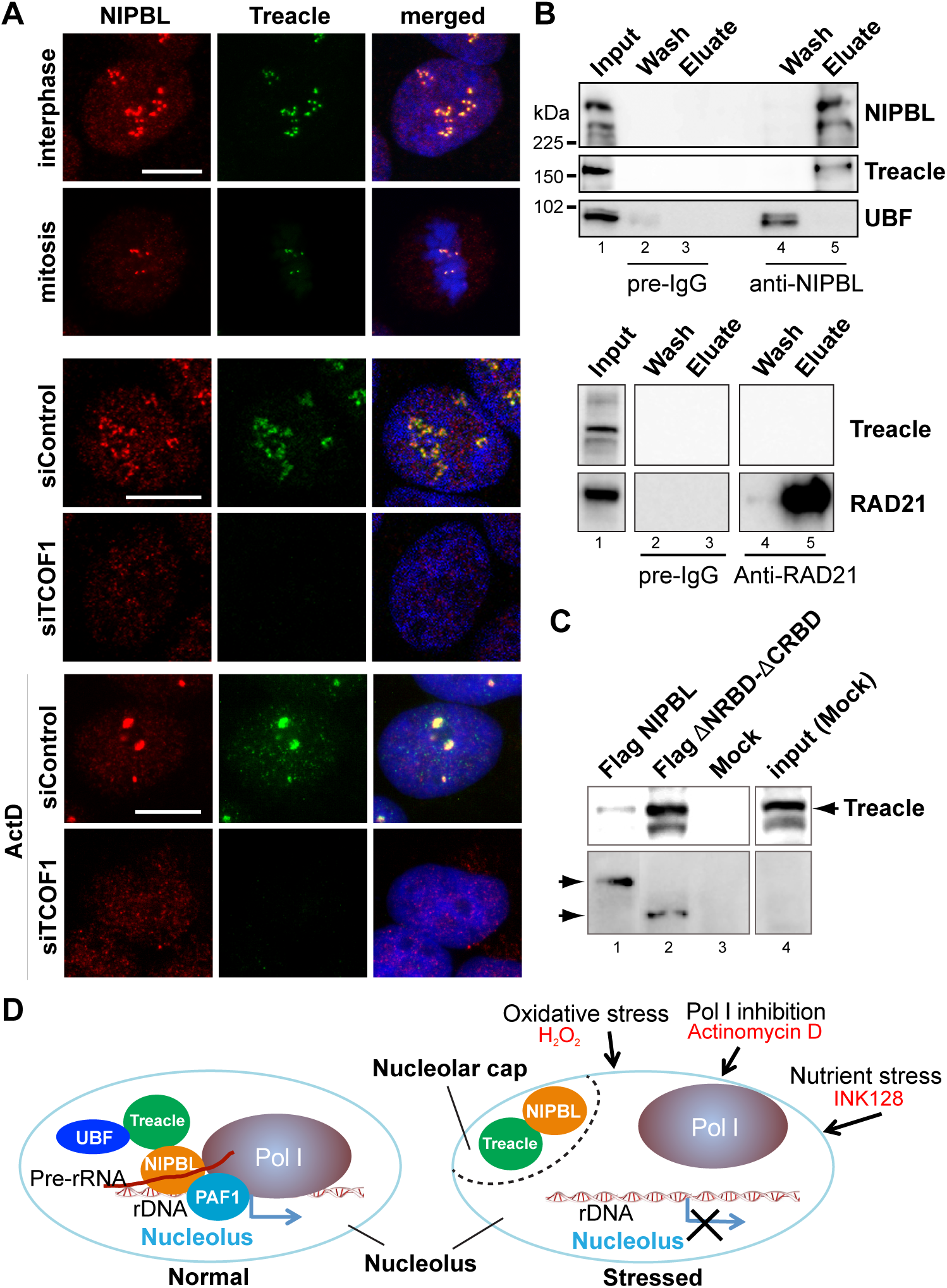
NIPBL interacts and closely colocalizes with treacle in the nucleolus. **A.** Top: Immunofluorescent staining of interphase and mitotic cells with antibodies specific for NIPBL and treacle as indicated. Middle and bottom: Treacle siRNA depletion abolishes NIPBL localization to the nucleolus and to the nucleolar cap structure with stress (induced by ActD treatment). **B.** Coimmunoprecipitation (co-IP)-western analysis the NIPBL interaction with treacle and UBF. Immunoprecipitation was performed using pre-immune IgG or anti-NIPBL antibody. Precipitates were washed with 1M KCl (Wash) and eluted with 2M guanidine-HCL (Eluate) (see Methods). Western blot was probed with antibodies specific for NIPBL, treacle and UBF as indicated. Similar co-IP-western using antibody specific for Rad21 that brings down the holo-cohesin complex probed with antibody specific for treacle, showing no signal (bottom). **C.** TCOF1 interaction is not affected by deletion of the NIPBL RNA binding domains. Mock-transfected cell extract or extracts from cells expressing FLAG-tagged full-length NIPBL or NIPBL mutant lacking NRBD and CRBD (ΔNRBD-ΔCRBD) were used for FLAG IP. Precipitated materials on beads were probed with antibody specific for TCOF1 (top) or FLAG (bottom). **D.** A schematic diagram of two-tiered recruitment and function of NIPBL in the nucleolus. Left: A subpopulation of NIPBL stably interacts with treacle that recruits NIPBL to the FC region in the nucleolus. Nascent pre-rRNA transcript further promotes NIPBL binding to rDNA chromatin, and binding of NIPBL to rDNA in turn recruits PAF1 and stimulates pre-rRNA synthesis. Right: Ribosomal RNA transcription is repressed by different types of stress. NIPBL dissociates from rDNA due to the reduced amount of pre-rRNA transcript, and is targeted to the nucleolar cap by treacle. Removal of NIPBL-treacle from rDNA further decreases rRNA production.

## Discussion

Our results demonstrate the novel RNA binding activity of NIPBL, which dictates its localization, rDNA binding and function in the nucleolus. Genome-wide analysis revealed the significant enrichment of NIPBL binding to 40S pre-rRNA. We identified two distinct RNA binding domains both required for efficient NIPBL binding to rRNA in vivo. Furthermore, treacle was identified to be the novel interaction partner for NIPBL, which directs NIPBL to the FC region during interphase, nucleolar organizer regions during mitosis, and nucleolar cap under stress conditions. At the rRNA loci, NIPBL promotes the recruitment of the PAF1 complex to facilitate rRNA transcription. Stress-induced inhibition of rRNA transcription facilitates the dissociation of NIPBL from rDNA loci and its sequestration into the nucleolar cap, further suppressing rRNA transcription. The NIPBL binding signal to the rDNA region is disproportionally strong compared to its binding to the unique cohesin binding sites, opposite from the cohesin binding signal, strongly suggesting that the rDNA loci are the major site of action for NIPBL, rather than cohesin.

### Multiple mechanisms of NIPBL targeting to chromatin

NIPBL loads cohesin onto chromatin. Although evidence suggests that cohesin interaction with sequence-specific binding proteins can determine the cohesin binding peaks (e.g., CTCF coinciding with cohesin peaks at the boundaries of topologically associated domains (Merkenschlager and Nora, 2016)), NIPBL localization also dictates cohesin binding regions in the genome. For example, NIPBL was shown to be recruited to the replication origins through the interactions with the pre-replication complex in Xenopus and human cells, which is critical for reloading of cohesin to the genome after its dissociation during cell division and establishment of sister chromatid cohesion on newly replicated DNA (Gillespie and Hirano, 2004; Takahashi et al., 2008; Takahashi et al., 2004; Zheng et al., 2018). NIPBL also loads cohesin onto the H3 lysine 9 methylation (H3K9me)-marked heterochromatin at both peri-centromeres and subtelomeres via heterochromatin binding protein HP1 homologs in yeast and human cells (Bernard et al., 2001; Dheur et al., 2011; Nonaka et al., 2002; Zeng et al., 2009a). NIPBL, and not cohesin, primarily interacts with HP1 (Fischer et al., 2009; Lechner et al., 2005; Serrano et al., 2009; Zeng et al., 2009a). In addition, NPBL interacts with the mediator, which promotes specific clustering of NIPBL/cohesin at the enhancer/promoter regions (Kagey et al., 2010). In the current study, we found another novel interacting partner treacle required for NIPBL targeting to the subnuclear organelle (nucleolus), and the critical role of the nascent RNA transcript in stable NIPBL binding to the rDNA loci. Thus, there are multiple mechanisms to direct targeting of NIPBL (and cohesin) to specific genomic regions to carry out their functions. Importantly, high salt-resistant stable binding of treacle-NIPBL interaction suggests that a subpopulation of NIPBL is stably engaged in this complex to be localized to the nucleolus. In contrast, cohesin failed to coprecipitate treacle. Furthermore, NIPBL binding appears to be more significant at the rDNA regions than at unique cohesin-binding sites. This is in contrast to cohesin binding at the unique cohesin binding sites, which is more pronounced than its binding at the rDNA regions. These results strongly suggest that the rDNA region is a major site of action for NIPBL.

### NIPBL in stress response

In addition to the interaction with treacle, we found specific binding of NIPBL to pre-ribosomal RNA species and RNA-dependent binding to ribosomal DNA loci. RNA transcript-dependent NIPBL recruitment suggests possible feedback mechanism. Consistently, stress-induced suppression of rRNA transcription resulted in relocalization of NIPBL from the FC region to the nucleolar periphery (nucleolar cap) or dispersion, and reduced binding to the rDNA region, which in turn would lead to further downregulation of rRNA transcription. Interestingly, SIRT7 also utilizes a similar rRNA-dependent recruitment mechanism to reinforce rRNA transcriptional repression in response to stress (Chen et al., 2013). The rRNA transcript-dependent recruitment mechanism of NIPBL, therefore, may be an effective way to ensure a timely silencing of rRNA transcription in response to environmental stress (Fig. 8D). It is interesting to note that the identified NIPBL domains containing RNA binding activities are not conserved in lower eukaryotes in contrast to the conserved cohesin and Scc4 interacting domains (Chao et al., 2015; Chao et al., 2017; Hinshaw et al., 2015; Kikuchi et al., 2016) (Fig. 5C). This raises the possibility that this represents a relatively newly acquired role during evolution.

### Multifaceted roles of NIPBL in ribosome biogenesis

Mutations of genes that impair rRNA transcription and ribosome biogenesis cause phenotypically related human disorders with growth and mental retardation, such as Treacher Collins Syndrome, Diamond-Blackfan anemia, and Shwachman–Diamond Syndrome, and are termed “ribosomopathies” (Hannan et al., 2012; Teng et al., 2013). Previous studies indicated that NIPBL/cohesin regulate many RNA Pol II genes involved in ribosome biogenesis and protein translation, and thus it was proposed that CdLS is also a type of ribosomopathy (Bose et al., 2012; Harris et al., 2014; Zakari et al., 2015). Indeed, L-leucine, which was found to be effective in treating ribosomopathies (Boultwood et al., 2013), can ameliorate certain developmental defects of Nipbl-depleted zebrafish (Baoshan Xu, 2014). Interestingly, NIPBL was shown to stimulate production of snoRNA, which guides chemical modification of rRNA critical for ribosome biogenesis (Zakari et al., 2015). Furthermore, Nipbl stimulates the expression of RNA processing genes, and Nipbl haploinsufficiency results in accumulation of unmodified RNAs activating the stress kinase PKR, which phosphorylates eIF2α and shuts down general protein translation (Yuen et al., 2016). In the current study, we found that NIPBL recruits the PAF1 complex and stimulates pre-ribosomal RNA synthesis. Since cohesin depletion also affects pol I transcription to a certain extent, we cannot determine whether the observed NIPBL function in pre-rRNA synthesis is entirely cohesin-independent. Nevertheless, these results together indicate that NIPBL mutation in CdLS may affect ribosome biogenesis and protein synthesis in three different ways: (1) decreased pre-rRNA synthesis, (2) decreased rRNA processing, and (3) decreased protein translation. Nucleolar dysfunction has been linked to both disrupted neurodevelopment and neurodegeneration (Hetman and Pietrzak, 2012; Narla and Ebert, 2010; Parlato and Kreiner, 2012). Thus, this aspect of NIPBL function may be critically linked to the growth and neurological phenotype of CdLS.

### Relationship between CdLS and TCS

The identified treacle interaction provides the first potentially direct link between CdLS and Treacher Collins syndrome (TCS) not inconsistent with the previous argument that CdLS is a type of ribosomopathy. Curiously, in Nipbl +/- mutant MEFs, *Tcof1* (encoding treacle) was shown to be downregulated (Kawauchi et al., 2009). It would be interesting to speculate that this treacle downregulation is to free up additional Nipbl for its canonical role in cohesin loading and chromosomal domain organization genome-wide to dampen the effect of Nipbl haploinsufficiency. It also raises a possibility that two genes (*TCOF1* and *NIPBL*) may serve as modifier genes in CdLS and TCS, respectively. While the physical link between NIPBL and treacle is provocative, the phenotype of TCS is highly restricted to facial bones and tissue with variable neuronal development phenotype (Kadakia et al., 2014; Sakai and Trainor, 2016; Yelick and Trainor, 2015), which differs from the complex multisystem phenotype of CdLS. The latter is most likely caused by multiple ways NIPBL affects nuclear functions, including cohesin-mediated and possibly cohesin-independent RNA Pol II transcriptional regulation in addition to the treacle-NIPBL pathway. Differential effects of NIPBL mutations on these processes in different cells/tissues may explain highly variable phenotype of the disease (Boyle et al., 2015; Mannini et al., 2013).

In conclusion, we identified a novel RNA binding activity of the essential cohesin-loading factor NIPBL, which is important for rRNA transcript-dependent binding of NIPBL to the rDNA region and regulation of rRNA transcription. It would be interesting to explore whether this transcript-dependent recruitment and transcriptional regulatory mechanism may be utilized at other NIPBL (and cohesin) target gene loci.

## Materials & Methods

### Cells and cell lines

Primary mouse embryonic fibroblasts (MEFs) derived from E15.5 wild type and Nipbl mutant embryos were described previously (Kawauchi et al., 2009). In summary, mice heterozygous for Nipbl mutation were generated (Nipbl +/-) from gene-trap-inserted ES cells. This mutation resulted in a net 30-50% decrease in Nipbl transcripts in the mice, along with many phenotypes characteristic of human CdLS patients (Kawauchi et al., 2009). Wild type and Nipbl +/- MEFs, immortalized wild type MEFs (Heale et al., 2006) and 293T cells were cultured at 37°C and 5% CO2 in DMEM (Gibco 31600-034) supplemented with 10% fetal bovine serum and penicillin-streptomycin (50U/mL).

### Antibodies

Rabbit polyclonal antibodies against NIPBL and MAU2 proteins were raised against bacterially expressed recombinant polypeptides and were antigen affinity-purified. Our NIPBL antibody targets the C terminus (a.a. 2430-2804) of human NIPBL variant A (NP_597677.2), and were used previously (Kong et al., 2014; Zeng et al., 2009a). This antibody was used previously for chromatin immunoprecipitation (ChIP) to demonstrate NIPBL/Nipbl binding at cohesin binding sites and its specificity was confirmed (Chien et al., 2011b; Zeng et al., 2009a). MAU2 antibody targets a.a. 125-467 of human MAU2 (NP_056144.3). Antigen affinity-purified rabbit polyclonal antibodies specific for Rad21, SMC1, and SMC3 and the preimmune IgG control were published previously (Gregson et al., 2001) and were used for chromatin immunoprecipitation (ChIP) as published (Parelho et al., 2008; Zeng et al., 2009b). Antibodies specific for B23 (GeneTex GTX10530), Fibrillarin (GeneTex GTX24566), UBF (Santa Cruz sc-13125X), Treacle (LifeSpan Bio LS-C198318 and PROTEINTECH GROUP, INC 11003-1-AP), CTCF (Millipore 07-729), α-tubulin (Sigma T9026), BrdU (GeneTex GTX27384), H3 (Abcam ab1791), and PAF1 (GeneTex GTX60361) were also used.

### Plasmid construction

To construct plasmid pGEX-6P-1-NIPBL-NRBD-N (a.a. 540-680), pGEX-6p1-NIPBL-NRBD-C (a.a. 681-919), NIPBL-NRBD-N and NIPBL-NRBD-C fragments were amplified from template plasmid pGEX-2TKN-Scc2-NRBD (a.a. 640-919) using primers 5’- AGCTAGGATCCGTTAGCATTGATCTTCATCA-3’, 5’-AGCTAGCGGCCGCTTATGTGTCAGACAGTCTATTTT-3’ and 5’-AGCTAGGATCCAAACCAAATGACAACAAACA-3’, 5’-AGCTAGCGGCCGCTTATTTGTCATCTTTACTAGTTG-3’, respectively. The amplified fragments were digested with BamH I and Not I then subcloned into the same sites of pGEX- 6P-1 vector (GE Healthcare). Halo-tagged NIPBL mutants “499-1124 a.a.” and “1844-2804 a.a.” (Oka et al., 2011) were digested with AsisI and PmeI and re-cloned into a modified pIRES vector containing N-terminal FLAG epitope. Two fragments NRBD and CRBD were amplified from plasmid Halo-tagged full length NIPBL (Oka et al., 2011) using two sets of primers (5’- ACAAGGACGACGATGACAAGGGATCCGCGATCGCCACAAAACCAAATGACAACAAAC-3’/ 5’-TGCGGCCGCTGCTGCAGCTGCTGCTTTGTCATCTTTACTAGTTGGTG-3’ AND 5’- GCAGCAGCTGCAGCAGCGGCCGCAAAGGACAAAAGGAAAGAGAGAAAATC-3’/5’- GTTATCTATGCGGCCGCCTAGAATTCAGTTTAAACGCTGGAAGTCCCATCCTTGGC-3’) which introduce a linker containing 8 alanine between them. Then these two fragments were assembled into the modified pIRES vector at AsisI and PmeI sites using Gibson Assembly kit (NEB) following manufacturer’s instructions. FLAG-tagged mutant “ΔNRBD-ΔCRBD” was obtained in the similar strategy by using another two sets of primers 5’-ACAAGGACGACGATGACAAGGGATCCGCGATCGCCATGAATGGGGATATGCCCCATGT- 3’/5’-TTGTTACCCTCTGTCCTTAATGCTGGCCTTGACCCATTTCCCG3’ and 5’-GGTCAAGGCCAGCATTAAGGACAGAGGGTAACAAGAGTAAAG-3’/5’- GTTATCTATGCGGCCGCCTAGAATTCAGTTTAAACTACCATAGACTCCTTGAATGACTGCAG-3’. For FLAG-tagged full length NIPBL, FLAG-tag coding sequences were introduced at the ATG start codon of the NIPBL gene in 293T cells by CRISPR/Cas9 genome editing (gRNA 5’- CCCCATTCATCCTGAATTTC-3’; phosphorothioate modified single-stranded oligonucleotide donor 5’-A*C*TTTTATAGGCAACACCATTCCAGAAATTCAGGATGGATTACAAGGATGACGATGACAA GAATGGGGATATGCCCCATGTCCCCATTACTACTC*T*T-3’) (Renaud et al., 2016). All plasmids and the FLAG-tagged NIPBL 293T cells were confirmed by DNA sequencing.

### Drugs used to induce nucleolar stress

All drugs were added to cell culture media and incubated for different lengths of time in the 37°C tissue culture incubator. Actinomycin D (Sigma) was used at a final concentration of 50 ng/ml and incubated for 2 hrs. Cells were treated with 0.5mM H_2_O_2_ (Ricca Chemical, 381916) for 3 hrs. mTOR inhibitor INK128 (Active Biochem, A-1023) was used at 100 nM and cells treated for 24 hrs. Alternatively, cells were treated with 3mM methyl methanosulfonate (MMS) for 1 hr, and fixed at 4 hr after MMS release.

### Immunofluorescence staining

MEFs and 293T cells were grown on coverslips in 24-well plates, fixed with 2% paraformaldehyde in PBS for 10 minutes, extracted with 0.2% Triton X-100 in PBS for 4 minutes, and blocked in PBS/ 2.5% BSA/ 5% heat-inactivated horse serum (Life Technologies 26050- 070)/ 5% goat serum (Life Technologies 16210-064) for 1 hr at room temperature. Primary and secondary antibodies were diluted in 1X PBS/ 1% BSA/ 5% heat-inactivated horse serum/ 5% goat serum. Coverslips were incubated in primary antibodies for 1 hr at room temperature followed by three PBS washes. Coverslips were incubated in secondary antibodies for 45 minutes at room temperature followed by three PBS washes. Then coverslips were stained with DAPI, washed with dH_2_O and mounted with Antifade (Life Technologies P-7481). For RNase-treated cell staining, MEFs grown on coverslips were first treated with 100ug/ml RNase A (Life Technologies 12091-021) for 10 min at room temperature and then stained using the protocol mentioned above. Staining in KD3 cells was done using a different protocol. Cells were fixed in 4% paraformaldehyde in PBS for 10 min, then blocked/ extracted in SNBP/ 0.1% Gelatin/ 4% heat-inactivated horse serum/ 4% goat serum/ 0.1% Triton-X for 30 min at 37°C. SNBP buffer is PBS with 0.02% saponin, 0.05% NaN_3_ and 1% BSA. Primary and secondary antibodies were diluted in SNBP/ 0.05% gelatin/ 1% heat-inactivated horse serum/ 1% goat serum. Coverslips were incubated in primary antibodies for 30 min at 37°C followed by three SNBP washes. Then coverslips were incubated in secondary antibodies for 30 min at 37°C followed by three SNBP washes. Then coverslips were stained with DAPI, washed with dH_2_O and mounted with Antifade.

### Nucleolar fractionation

Nucleolar fractionation on MEFs was done according to the following protocol (http://www.lamondlab.com/pdf/noprotocol.pdf) with some modifications. 5×10^7^ cells were used for each nucleolar fractionation. All buffers (S1, S2 and S3) were reduced to half except for buffer A (5ml). Five ml of cytoplasmic extract and three ml of nuclear extract were obtained from 5×10^7^ cells. The pelleted nucleoli were resuspended in 100ul buffer S2. Same volume of each fraction was loaded onto SDS gel for further analysis.

### Coimmunoprecipitation

Nuclear extract preparation and coimmunoprecipitation (co-IP) were performed as described previously (Heale et al., 2006). Briefly, Briefly, the antibody-protein complex was washed four times with HEMG buffer (25mM <u>H</u>EPES [pH 7.6], 0.1mM <u>E</u>DTA 12.5mM <u>M</u>gCl_2_, 10% <u>g</u>lycerol) containing 0.1M KCl and 0.1% Nonidet P-40. Bound proteins were eluted from the beads with HEMG-1.0M KCl (1.0M KCl) (“Wash”) followed by 2M guanidine-HCl (“Eluate”). Eluted proteins were precipitated with trichloroacetic acid (TCA) and analyzed by SDS-PAGE and western blotting.

### siRNA depletion

MEF cells were transfected using HiPerFect (Qiagen) following the manufacturer’s protocol. Media with siRNA and transfection reagents were removed 6 hours post-transfection and fresh media were added. Second round of transfection was performed 24 hours after first transfection. Cells were harvested 48 - 72 hours after the first transfection. Mouse Nipbl siRNAs (Nipbl-1: 5’- GTGGTCGTTACCGAAACCGAA-3’; Nipbl-2: 5’-AAGGCAGTACTTAGACTTTAA-3’), Rad21 siRNA (5’-CTCGAGAATGGTAATTGTATA-3’), human Treacle siRNA (5’- AGACTAGCATCTACCAACT-3’) (Lin and Yeh, 2009), control siRNA (5’- AATTCTCCGAACGTGTCACGT-3’) as well as human NIPBL siRNA (5’-CTAGCTGACTCTGACAATAAA-3’) were used. For mouse CTCF depletion, Dharmacon siGENOME Mouse CTCF (13018) siRNA - SMARTpool (Cat# M-044693-01-0010) was used.

### UV cross-linking and immunoprecipitation

UV cross-linking and immunoprecipitation (CLIP) was done according to previous studies with some modifications (Konig et al., 2010; Yao et al., 2012). RNA labeling was performed as described in the individual nucleotide CLIP sequencing (iCLIPseq) previously published (Konig et al., 2011). The modifications are described as the following. After cell lysis, 15 µl RQ1 DNase (Promega M610A) and 2 µl RNaseOut (Life Technologies 10777-019) were added to the lysate and incubate at 37°C for 3 min. RNaseOut was added to the lysate in order to maximize the recovery of intact RNAs. Approximately 5 µg of antibody was used in each IP. For CLIP followed by RT-qPCR, 2×10^6^ cells and 50 µg of Protein A Dynabeads (Life Technologies) were used per IP. After IP overnight and the washes, beads were resuspended in 25 µl RNase-free water. Five min incubation at 95°C was done with gentle shaking. Then the 25 µl supernatants were divided into two tubes for reverse transcription using the protocol of SuperScript II reverse transcription (Life Technologies). Reverse transcriptase was added into one tube (RT+) but not the other (RT-). For the comparison of the CLIP signals between the two treatments (e. g., control and Nipbl siRNA-treated cells), 20% of the cell samples were taken as input and subjected to total RNA extraction. The same volumes of eluted total RNA were taken for reverse transcription. The IP signals from different treatments were normalized to the input during data analyses. For CLIP-SDS-PAGE, 10^7^ cells and 100 µl Protein A beads slurry were used per IP. Precipitated RNA 3’ends were first dephosphorylated by shrimp alkaline phosphatase (SAP) (NEB M0371), and 3’ RNA linker (20 µM) (Dharmacon, 5’-phosphate-AGAUCGGAAGAGCGGUUCAG-3’ was added by T4 RNA Ligase 1 (NEB M0204) at 16°C overnight. The 5’ ends were radiolabeled with P^32^-γ-ATP (PerkinElmer) by T4 polynucleotide kinase (NEB M0201) at 37°C for 10 min. The precipitated and radiolabeled protein-RNA complexes on beads were heat-denatured and loaded on a 4-12% NuPAGE Bis-Tris gel (ThermoFisher NP0321BOX) with 1X MOPS running buffer (ThermoFisher NP0001). The protein-RNA complexes were transferred to a nitrocellulose membrane and exposed to the phosphorimager screen.

### CLIP-sequencing

eCLIP of NIPBL in K562 cells was performed as previously described, including standard read processing and processing of reads which uniquely mapped to the genome (Van Nostrand et al., 2016), with the exception of the use of NuPAGE Tris-Acetate (3-8%, ThermoFisher) protein gels to ensure proper separation of the large NIPBL protein. Quantification of ribosomal RNA binding was performed using a family-aware repeat element mapping pipeline to identify reads that only mapped to 45S pre-rRNA, 18S rRNA, or 28S rRNA respectively (Van Nostrand, E.L., *et al. in preparation*). To normalize relative enrichment between IP and input, relative information was calculated as the Kullback-Leibler divergence (relative entropy): 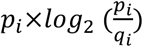, where *p_i_* is the fraction of total reads in IP that map to a queried element *i* (peak, gene, or repetitive element), and *q_i_* is the fraction of total reads in input for the same element. NIPBL eCLIP data has been deposited at the ENCODE Data Coordination Center webportal as KY_SCC2C_01 (lot # 093015) (https://www.encodeproject.org/).

### In vitro RNA pull-down using GST fusion proteins

Purifications of GST-tagged proteins were described previously (Hisaoka et al., 2014). GST and GST-tagged proteins were incubated with glutathione-Sepharose beads in HEMG binding buffer (25 mM Hepes-KOH pH 7.9, 0.1 mM EDTA, 12.5 mM MgCl2, 20% vol/vol glycerol, and 1 mM DTT, 0.1M KCl, 0.1% Nonidet P-40) (for experiments in Figs. 3c, 3d and S3b) or binding buffer A (0.1% Triton X-100, 50 mM Tris-HCl pH7.9, 0.5 mM PMSF, 150 mM NaCl) (Figs. 3e, 3f and S3a) for 45 min, the unbound proteins were then removed by washing with buffer A twice. The beads immobilized proteins were mixed with total RNA (10 µg) and rotated at 4°C for 1 h followed by extensive washing with the same buffer. Proteins were then digested by proteinase K and associated RNAs were purified by phenol-chloroform extraction and ethanol precipitation. The purified RNA was separated by formaldehyde-agarose gel stained by ethidium bromide, urea-PAGE stained by GelRed (Biotium) and was also used for RT-qPCR. For RT-qPCR analysis of bound RNA species, cDNA was prepared from purified total RNA or RNA co-purified with GST pull-down using ReverTraAce (Toyobo) with random primers. Real-time PCR was carried out in triplicate with primers by using SYBR Green Realtime PCR Master Mix-Plus (Toyobo) in the Thermal Cycler Dice Real-Time PCR system (TaKaRa). PCR primers are listed in Supplemental Tables S2 and S3.

### ChIP with RNase treatment

Approximately 6 × 10^6^ cells were used per IP. Cells were cross-linked with 1% formaldehyde for 10 min at room temperature. Glycine was added to a final concentration of 0.125 M to stop cross-linking. Cells were washed twice with PBS and collected by scraping. Approximately 2 × 10^7^ cells were resuspended in 1ml of Farnham lysis buffer (5 mM PIPES pH 8.0 / 85 mM KCl / 0.5% NP-40/ protease Inhibitors), centrifuged for 5min 2000rpm at 4°C. In terms of RNase-treated ChIP, pellets were re-suspended in 1 ml RIPA buffer (PBS / 1% NP-40 / 0.5% sodium deoxycholate / 0.1% SDS/ protease Inhibitors). RNase A was added to the lysate with final concentration of 500ug/ml. Both the lysates with and without RNase A were incubated at 37°C for 1 hour. SDS was added to the lysate after RNase treatment to a final concentration of 0.5% to facilitate sonication. For other ChIP experiments in the study, sample pellets were resuspended in SDS buffer (50mM Tris-HCl pH 8.0/ 10mM EDTA/ 1% SDS) after Farnham lysis and subjected to sonication. The lysates were sonicated using Bioruptor (Diagenode UCD-200) to fragments averaging around 300-500bp. The extracts were diluted with ChIP dilution buffer (0.01% SDS, 1.1% Triton X-100, 1.2 mM EDTA, 16.7 mM Tris-HCl (pH 8.1), 167 mM NaCl) with protease inhibitors and centrifuged for 10 min at 10,000 g at 4°C. Extracts were precleared for 1 h with protein A-Sepharose (GE Healthcare) supplemented with 1mg/ml BSA. Consistently, 10% of the extract for each sample was taken as input DNA. Antibodies was added to the extracts for each IP and incubated overnight at 4°C on a rotator platform. The amount of antibody used in ChIP is documented in Table 3-2. The next day, the antibody-bound complexes were immunoprecipitated with Protein A-Sepharose beads for 1 h and subsequently washed with low-salt buffer (0.1% SDS/ 1% Triton X-100/ 2 mM EDTA/ 20 mM Tris-HCl pH 8/ 150 mM NaCl), high-salt buffer (0.1% SDS/ 1% Triton X-100/ 2 mM EDTA, 20 mM Tris-HCl pH 8/ 500 mM NaCl), lithium salt buffer (0.25 M LiCl/ 1% Nonidet P-40/ 1% deoxycholate/ 1 mM EDTA/ 10mM Tris-HCl pH 8), and TE (10mM Tris-HCl/ 1 mM EDTA pH 8.0). DNA was eluted off the beads with 200ul of elution buffer (1% SDS/ 0.1 M NaHCO3) on a rotator platform at room temperature for 1 hour. The eluates were collected and crosslinking was reversed overnight at 65°C along with the input lysates. Quantitative PCR (q-PCR) was performed using the CFX96 real-time PCR detection system (Bio-Rad) with SYBR premix Ex Taq II (Clontech RR820B). ChIP DNA was purified with Qiagen PCR purification kits. ChIP signal was normalized by subtracting the preimmune IgG ChIP signal, then divided by input DNA signal. Primers used are listed in Supplemental Table S2.

### RT-qPCR

Total RNA was extracted using the Qiagen RNeasy Plus kit. First-strand cDNA synthesis was performed with SuperScript II reverse transcriptase (Life Technologies). Quantitative PCR (q- PCR) was performed using the CFX96 real-time PCR detection system (Bio-Rad) with SYBR premix Ex Taq II (Clontech RR820B). Standard curves were generated for each primer pair using a serial dilution of cDNA. Values were generated based on threshold cycles (Ct) with respect to the efficiency of each primer pair. Pre-rRNA expression is normalized to *Rnh1* (house-keeping gene) expression. Primers used are listed in Supplemental Tables S2 and S3.

### FU labeling

5-fluorouridine (Sigma, F5130) was added to cell culture with final concentration of 2mM, followed by 5 min incubation in cell culture incubator. After 5 min, the 5-fluorouridine-containing medium was removed and the cells were kept in growth media for 30 min in the cell culture incubator. After 30 min, cells were fixed and subjected to immunofluorescence staining.

### Statistical Analyses

All collected data were included in the analyses. At least three biological replicates per condition were used for all the experiments. A biological replicate is defined as an independent culture of cells that was separately manipulated and subsequently analyzed. For ChIP, CLIP, RNA pull-down and RT-qPCR experiments, data was analyzed using two-tailed Student’s t-test.

## Acknowledgement

This work was supported in part by National Institutes of Health (HG009530 to E.V.N, NS075449, HG007005, and HG004659 to G.W.Y, and HD078849 to K.Y.).

---

**Supplemental Table S1.**
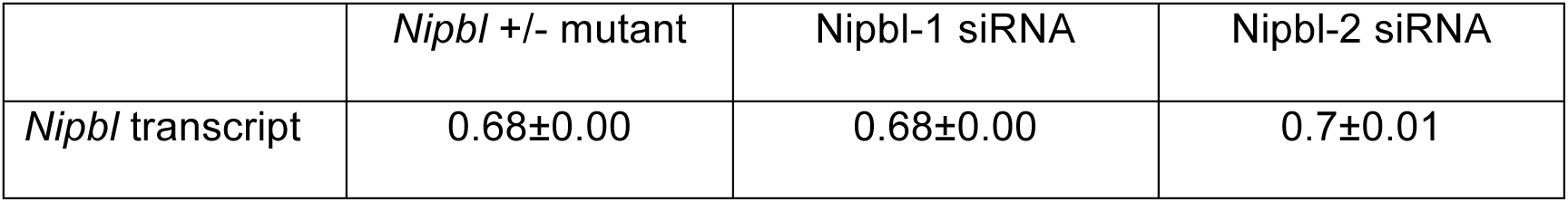
*Nipbl* transcript depletion.

**Supplemental Table S2.**
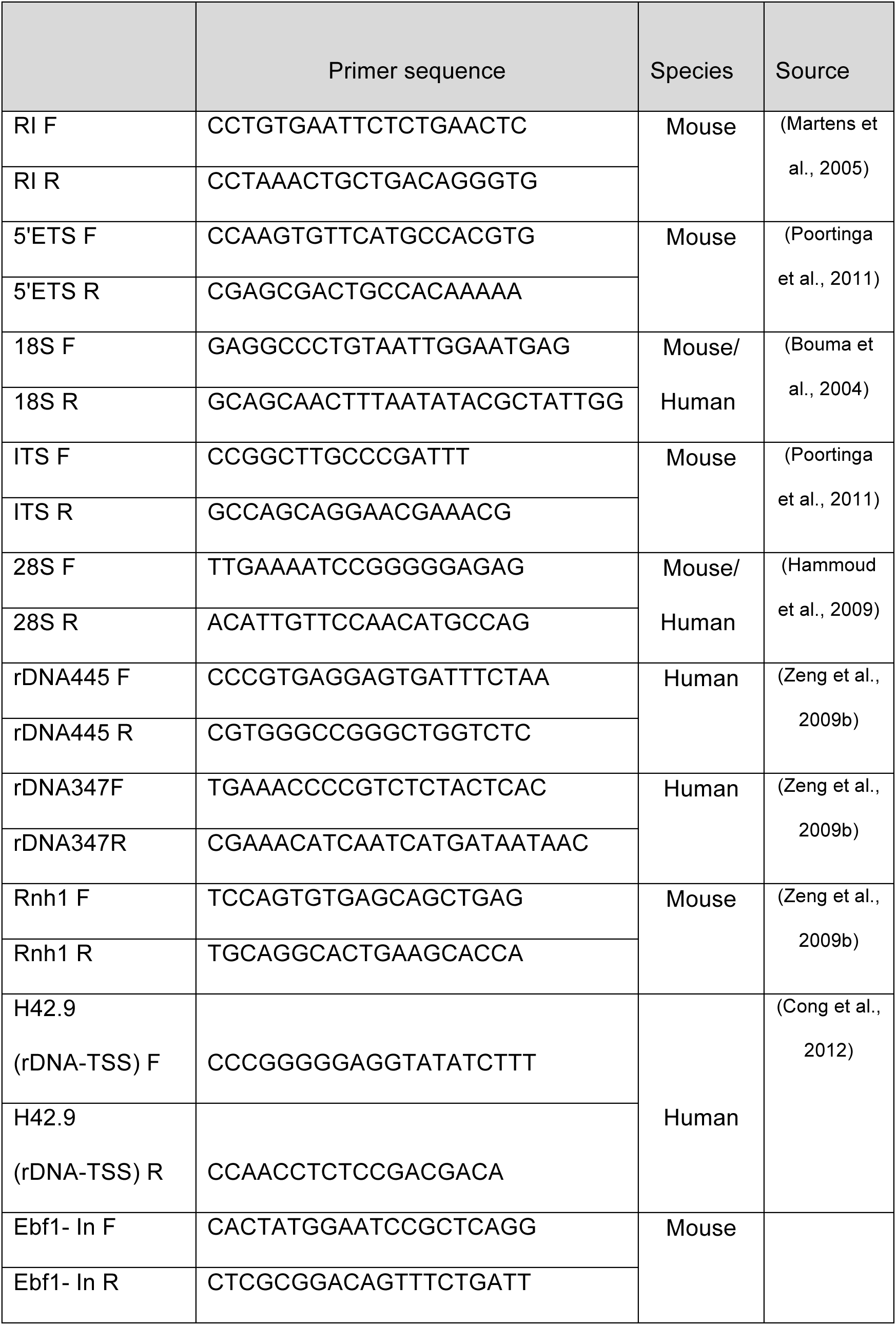
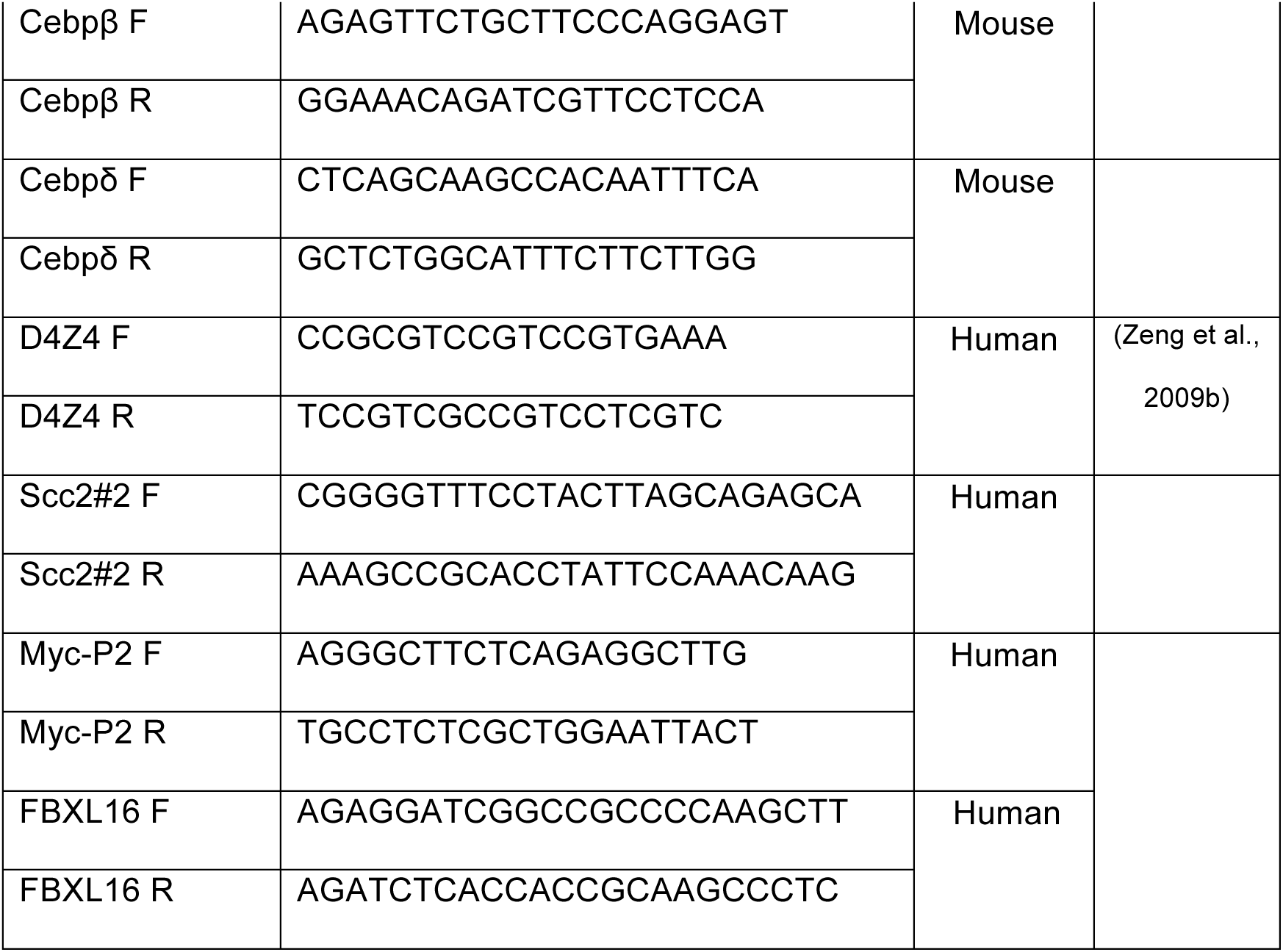
List of PCR primers.

**Supplemental Table S3.**
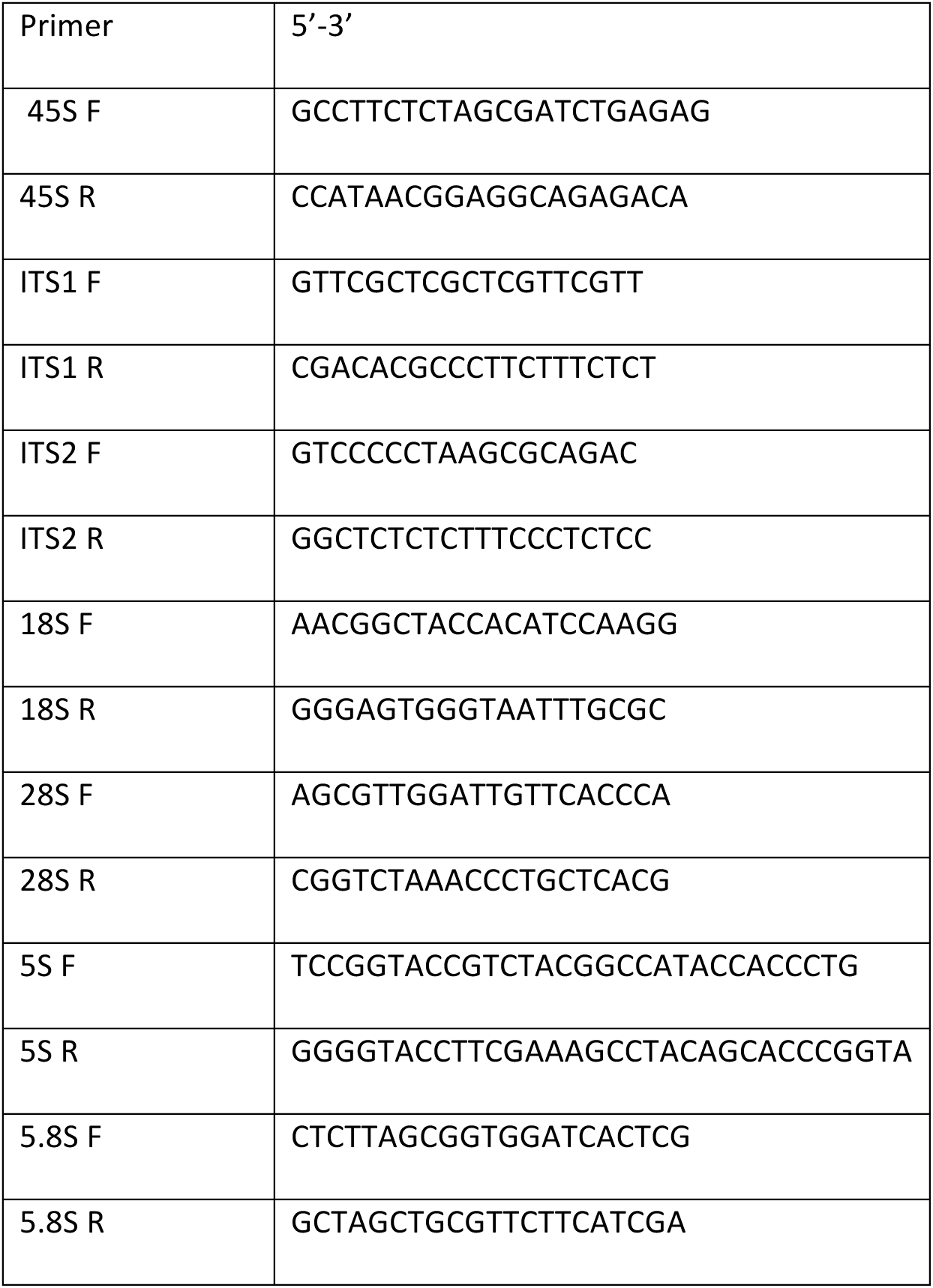
List of additional PCR primers (for Supplemental Figure S3a)

**Supplemental Fig. S1.**
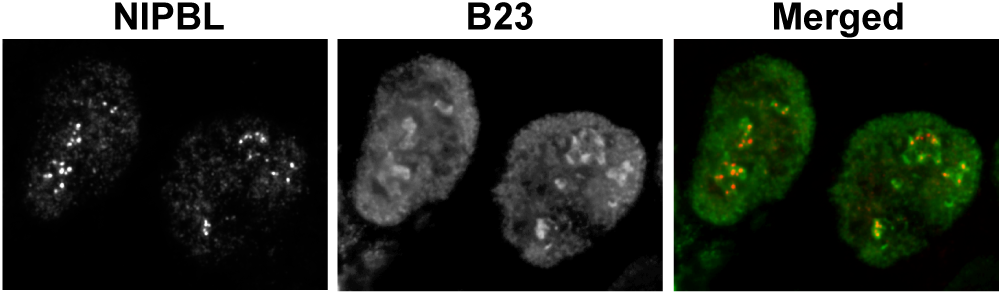
Localization of human NIPBL in the nucleolus. 293T cells are stained with antibody specific for NIPBL and B23.

**Supplemental Fig. S2.**
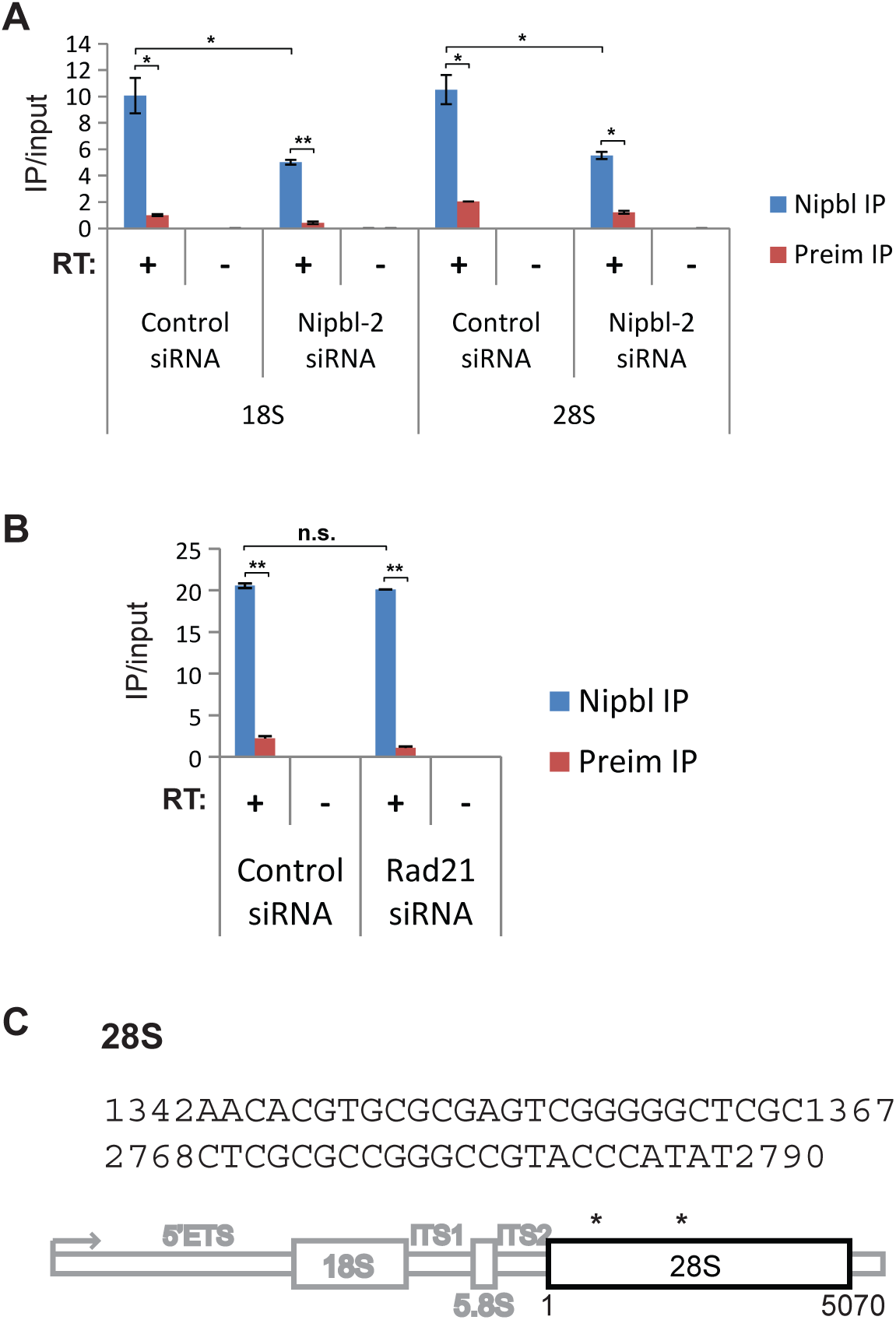
Nipbl binding to the coding regions of rRNA. **A.** The second siRNA specific for Nipbl reduces Nipbl binding to 18S and 28S rRNAs. Depletion efficiency is shown in Supplemental Table S1. **B.** Rad21 depletion has no effect on Nipbl binding to rRNA. Nipbl ChIP-qPCR using 18S primers was performed in control siRNA- or Rad21 siRNA-treated cells. For both (A) and (B), data is the mean ± SD of three biological replicates. * p<=0.05; ** p<=0.01; ns, not significant, p>0.05. **C.** 28S sequences recovered from CLIP RNA cloning and sequencing analysis. The region above 250kDa (indicated by the bracket in Figure 2C) was excised from the gel and isolated RNA was reverse-transcribed and cloned into a plasmid and sequenced. The locations of two RNA sequences are shown below.

**Supplemental Fig. S3.**
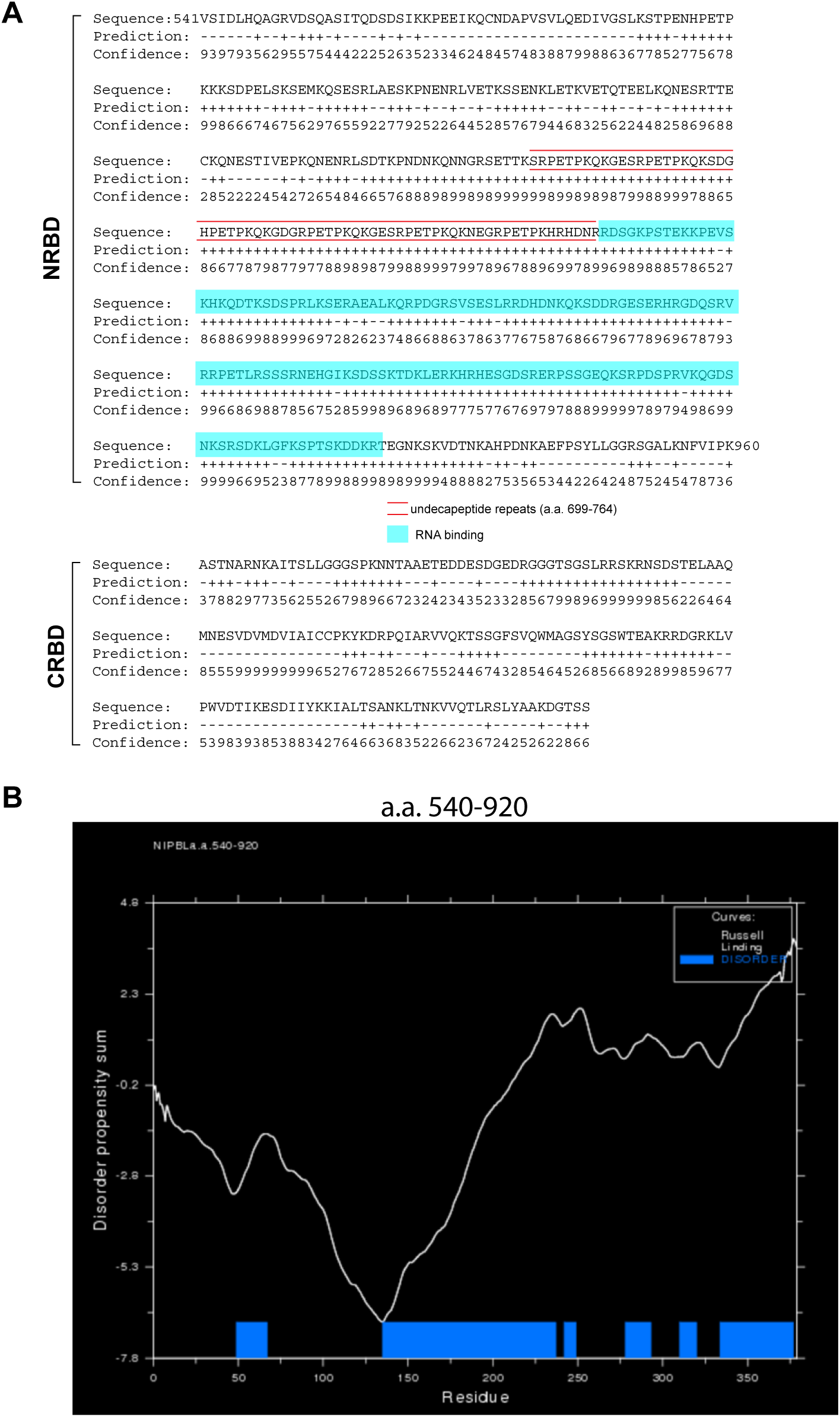
NRBD contains disordered domains. **A.** BindN analysis of NIPBL NRBD and CRBD. NRBD located downstream of the undecapeptide repeats as indicated. **B.** Analysis of the NRBD (a.a. 540-919) by Globeplot 2 (Intrinsic Protein Disorder, Domain & Globularity Prediction) (http://globplot.embl.de).

**Supplemental Fig. S4.**
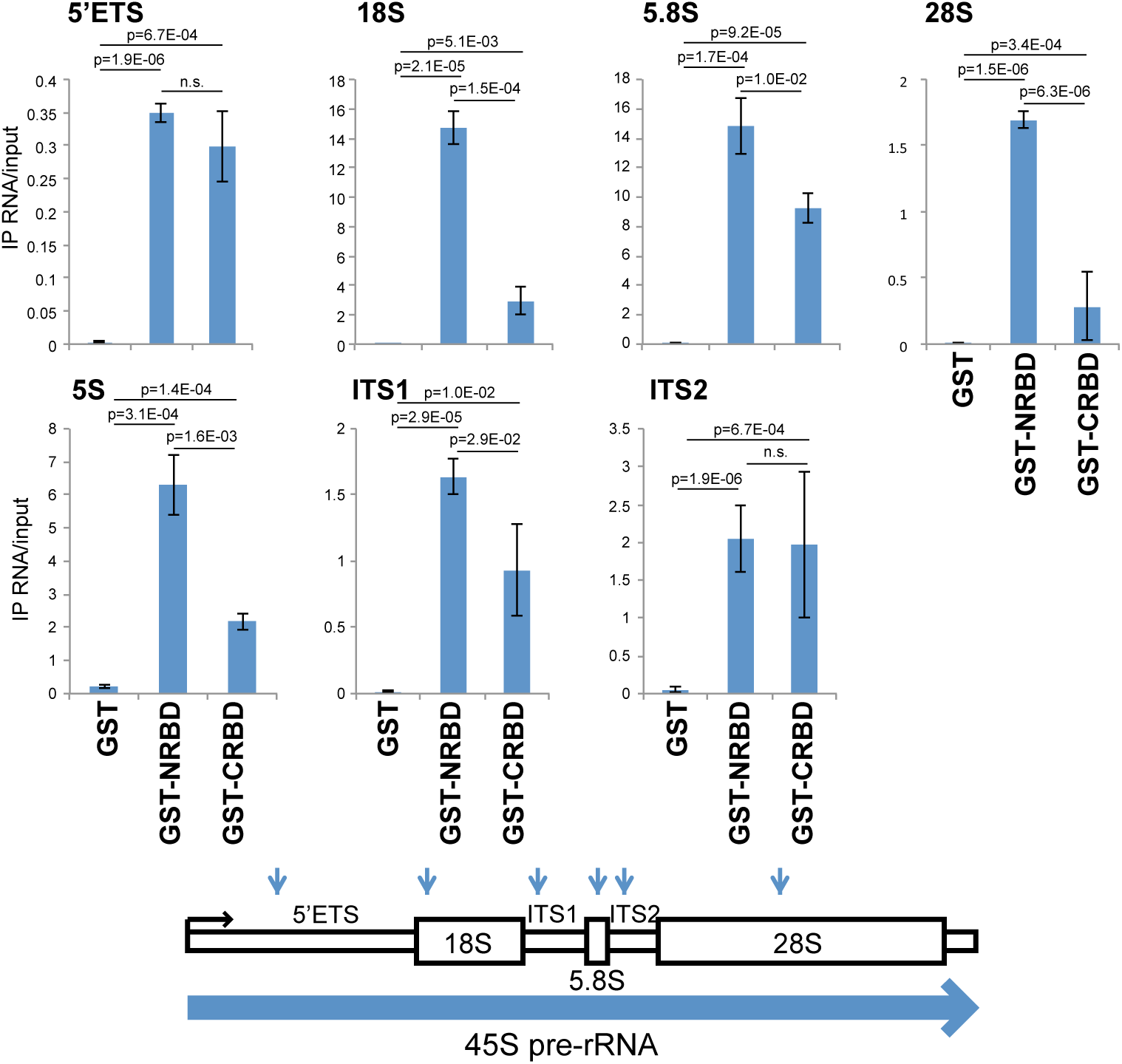
In vitro rRNA binding analysis of Nipbl RNA binding domains. In vitro rRNA binding to GST, GST-NRBD and GST-CRBD was analyzed as in Figure 4C, using an additional set of PCR primers against rRNA (Supplemental Table S2) and binding buffer A (0.1% Triton X-100, 50 mM Tris-HCl pH7.9, 0.5 mM PMSF, 150 mM NaCl). Locations of PCR products are shown in Supplemental Table S3. Data is the mean ± SD of three biological replicates. GST-GFP is used as a negative control. n.s.: not significant.

**Supplemental Fig. S5.**
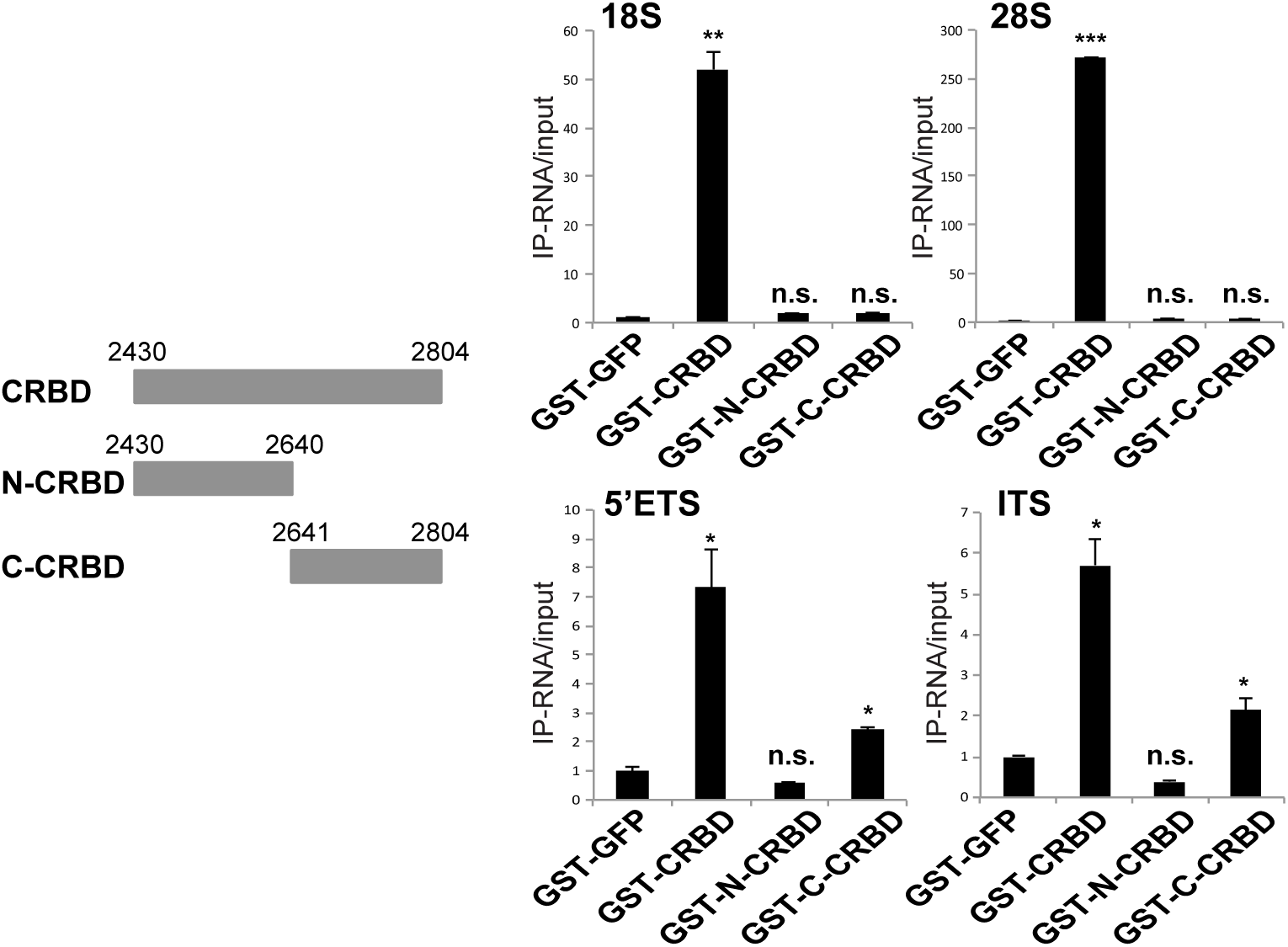
Division of CRBD into two fragments abolished their binding to 18S and 28S. GST-GFP (negative control), GST-CRBD, GST-N-CRBD (the N-terminal half of CRBD), and GST-C-CRBD (the C-terminal half of CRBD) as indicated were compared. Interestingly, C-CRBD retained some RNA binding activity towards 5’ETS and ITS regions. Data is the mean ± SD of three biological replicates, where statistical significance was calculated by comparing GST-tagged GFP to GST-tagged NIPBL deletion mutants at indicated region using a Student’s two-tailed t-test. * p<=0.05; ** p<=0.01; *** p<=0.001; ns, not significant, p>0.05.

**Supplemental Fig. S6.**
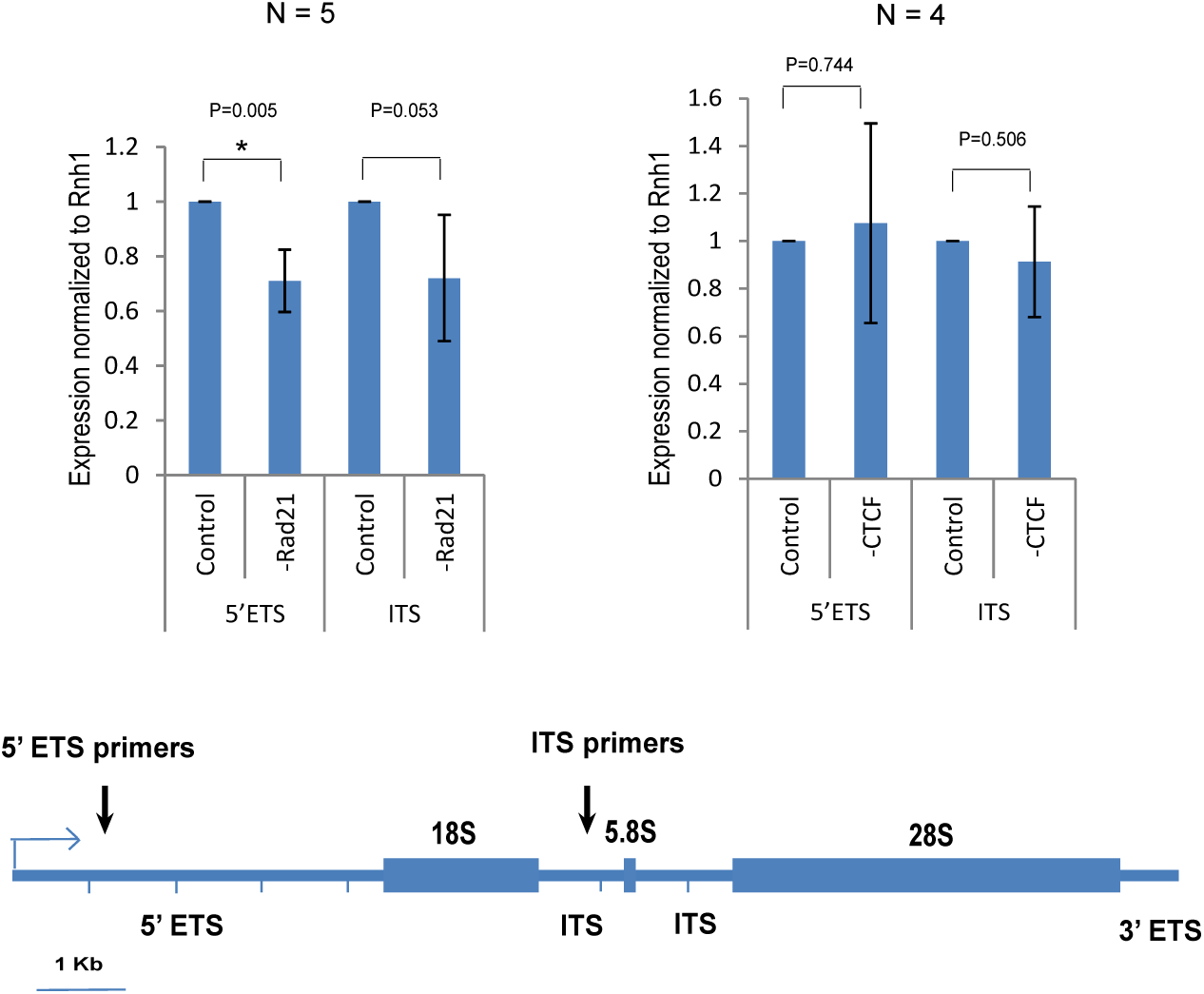
The effects of Rad21 or CTCF depletion on pre-rRNA synthesis. Experiments are performed as in Figure 6C. Rad21 (cohesin) depletion resulted in a modest decrease of pre-rRNA (as detected by the PCR primers specific for 5’ETS and ITS as indicated below) while CTCF depletion showed no significant effect. Data is the mean ± SD of three biological replicates, where statistical significance was calculated by comparing specific knockdown to control siRNA treatment at each region using a Student’s two-tailed t-test.

**Supplemental Fig. S7.**
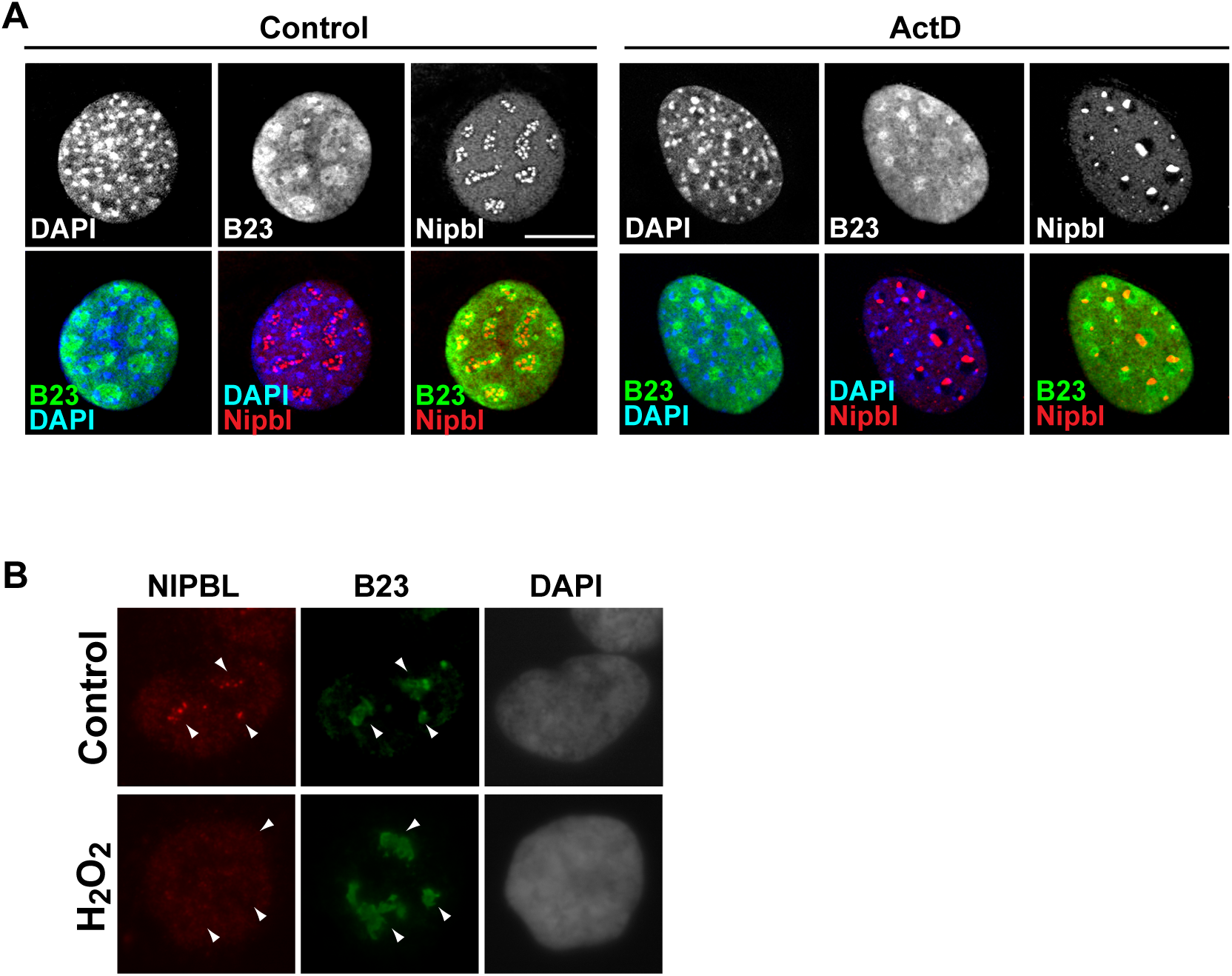
Nipbl delocalization upon stress induction. **A.** Actinomycin D (Act D) treatment as in Figure 5a co-stained with anti-B23 antibody. Upon treatment, Nipbl foci clusters to one side of the nucleoli. **B.** H_2_O_2_ treatment of human myoblasts resulted in displacement of NIPBL from the nucleolus.

**Supplemental Fig. S8.**
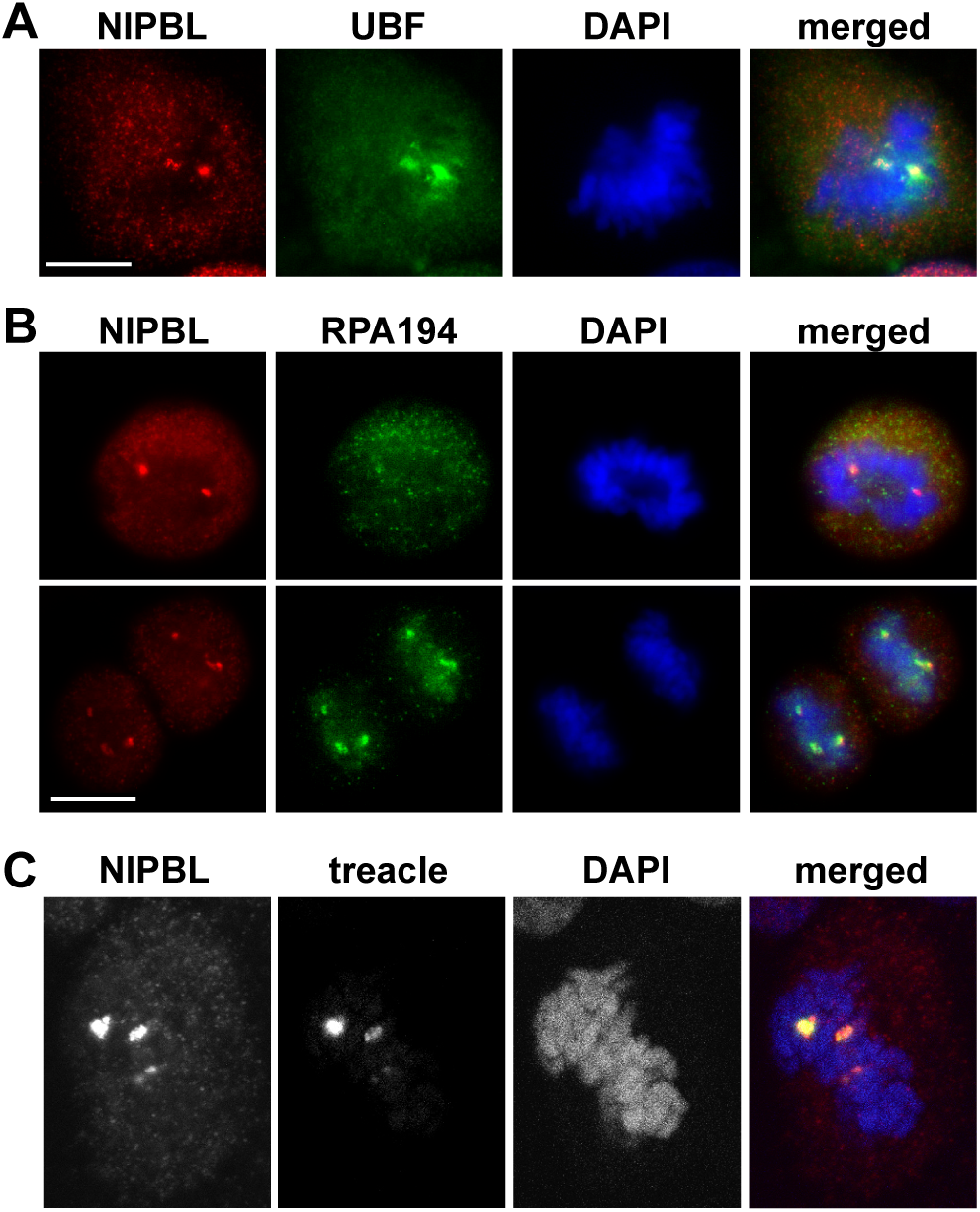
NIPBL localization at the NOC in mitosis. Immunofluorescent staining of NIPBL costained with antibodies specific for UBF **(A)**, Pol I subunit RPA194 **(B)**, and treacle **(C)**, demonstrating their colocalization at NOC.

**Supplemental Fig. S9.**
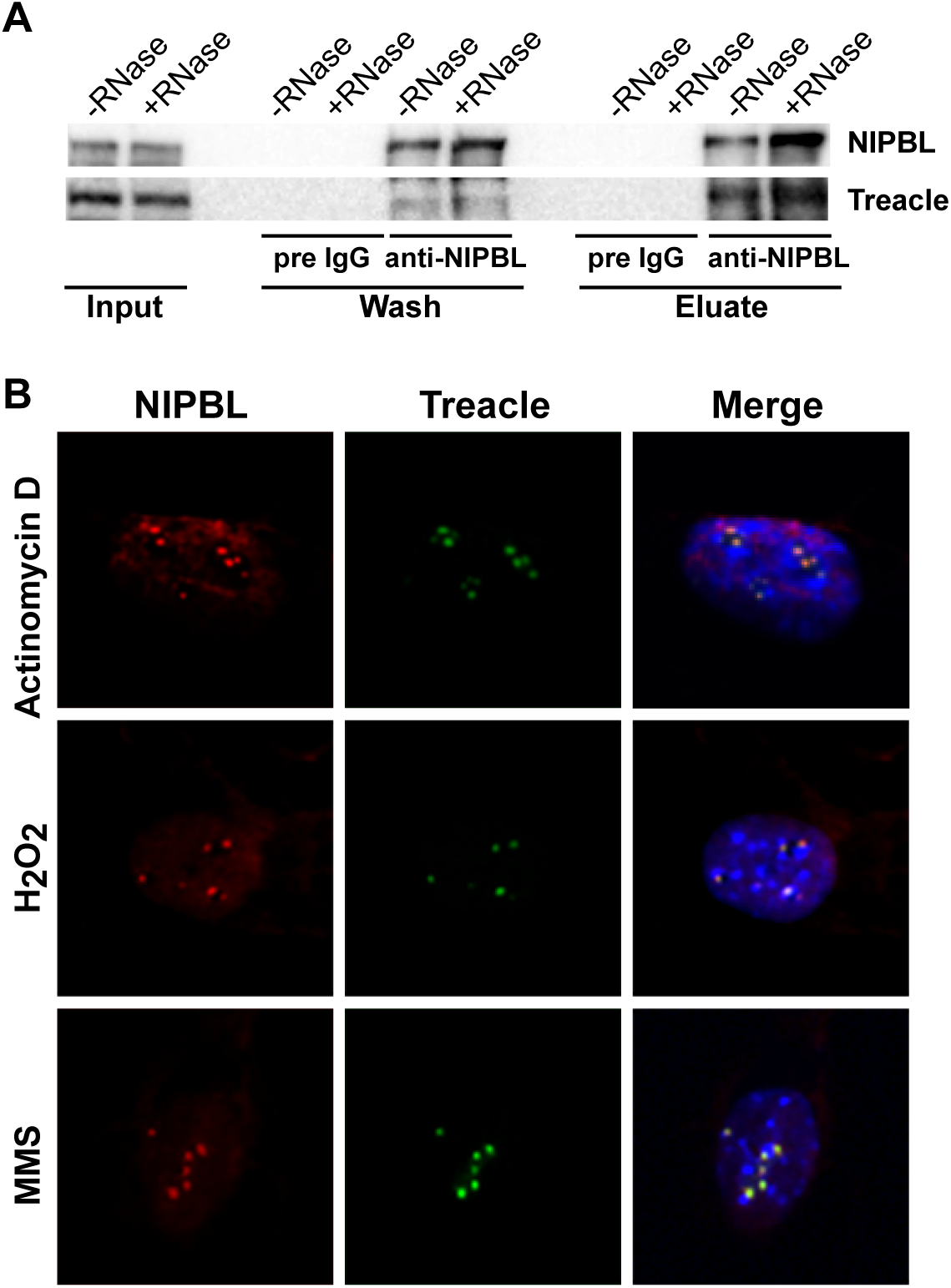
Interaction and colocalization of NIPBL and treacle. **A.** The NIPBL-treacle interaction is not affected by RNase treatment. **B.** Immunostaining of cells treated with ActD, H2O2 or MMS using antibody specific for NIPBL or treacle as indicated.

**Supplemental Fig. S10.**
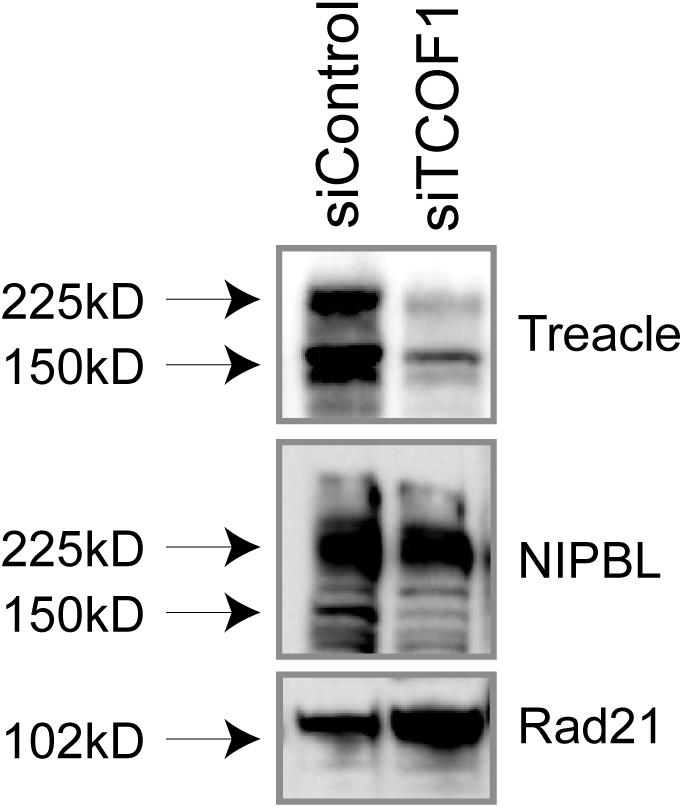
Western blot analysis of treacle depletion by TCOF1 siRNA. The nuclear extracts of 293T cells transfected with either control or TCOF1 siRNA were probed with antibodies specific for treacle, NIPBL and Rad21 as indicated.

## References

Baoshan Xu, N.S.a.J.G. (2014). L-Leucine partially ameliorates developmental deficits in zebrafish models for Cornelia de Lange syndrome. The FASEB Journal 28.

Bernard, P., Maure, J.-F., Partridge, J.F., Genier, S., Javerzat, J.-P., and Allshire, R.C. (2001). Requirement of heterochromatin for cohesion at centromeres. Science 294, 2539–2542.

Boisvert, F.M., Hendzel, M.J., and Bazett-Jones, D.P. (2000). Promyelocytic leukemia (PML) nuclear bodies are protein structures that do not accumulate RNA. J Cell Biol 148, 283–292.

Bose, T., Lee, K.K., Lu, S., Xu, B., Harris, B., Slaughter, B., Unruh, J., Garrett, A., McDowell, W., Box, A., et al. (2012). Cohesin proteins promote ribosomal RNA production and protein translation in yeast and human cells. PLoS Genet 8, e1002749.

Boulon, S., Westman, B.J., Hutten, S., Boisvert, F.M., and Lamond, A.I. (2010a). The nucleolus under stress. Molecular cell 40, 216–227.

Boulon, S., Westman, B.J., Hutten, S., Boisvert, F.M., and Lamond, A.I. (2010b). The nucleolus under stress. Mol Cell 40, 216–227.

Boultwood, J., Yip, B.H., Vuppusetty, C., Pellagatti, A., and Wainscoat, J.S. (2013). Activation of the mTOR pathway by the amino acid (L)-leucine in the 5q-syndrome and other ribosomopathies. Adv Biol Regul 53, 8–17.

Bouma, G.J., Hart, G.T., Washburn, L.L., Recknagel, A.K., and Eicher, E.M. (2004). Using real time RT-PCR analysis to determine multiple gene expression patterns during XX and XY mouse fetal gonad development. Gene Expr Patterns 5, 141–149.

Boyle, M.I., Jespersgaard, C., Brøndum-Nielsen, K., Bisgaard, A.M., and Tümer, Z. (2015). Cornelia de Lange syndrome. Clin Genet 88, 1–12.

Calabretta, S., and Richard, S. (2015). Emerging Roles of Disordered Sequences in RNA-Binding Proteins. Trends Biochem Sci 40, 662–672.

Chan, S.L., Huppertz, I., Yao, C., Weng, L., Moresco, J.J., Yates, J.r., Ule, J., Manley, J.L., and Shi, Y. (2014). CPSF30 and Wdr33 directly bind to AAUAAA in mammalian mRNA 3’ processing. Genes Dev 28, 2370–2380.

Chao, W.C., Murayama, Y., Muñoz, S., Costa, A., Uhlmann, F., and Singleton, M.R. (2015). Structural Studies Reveal the Functional Modularity of the Scc2-Scc4 Cohesin Loader. Cell Rep 12, 719–725.

Chao, W.C., Murayama, Y., Muñoz, S., Jones, A.W., Wade, B.O., Purkiss, A.G., Hu, X.W., Borg, A., Snijders, A.P., Uhlmann, F., et al. (2017). Structure of the cohesin loader Scc2. Nat Commun 8, 13952.

Chen, S., Seiler, J., Santiago-Reichelt, M., Felbel, K., Grummt, I., and Voit, R. (2013). Repression of RNA polymerase I upon stress is caused by inhibition of RNA-dependent deacetylation of PAF53 by SIRT7. Molecular cell 52, 303–313.

Chien, R., Zeng, W., Ball, A.R., and Yokomori, K. (2011a). Cohesin: a critical chromatin organizer in mammalian gene regulation. Biochem Cell Biol 89, 445–458.

Chien, R., Zeng, W., Kawauchi, S., Bender, M.A., Santos, R., Gregson, H.C., Schmiesing, J.A., Newkirk, D., Kong, X., Ball, A.R.J., et al. (2011b). Cohesin mediates chromatin interactions that regulate mammalian β-globin expression. J Biol Chem 286, 17870–17878.

Cong, R., Das, S., Ugrinova, I., Kumar, S., Mongelard, F., Wong, J., and Bouvet, P. (2012). Interaction of nucleolin with ribosomal RNA genes and its role in RNA polymerase I transcription. Nucleic acids research 40, 9441–9454.

Deardorff, M.A., Kaur, M., Yaeger, D., Rampuria, A., Korolev, S., Pie, J., Gil-Rodríguez, C., Arnedo, M., Loeys, B., Kline, A.D., et al. (2007). Mutations in cohesin complex members SMC3 and SMC1A cause a mild variant of cornelia de Lange syndrome with predominant mental retardation. Am J Hum Genet 80, 485–494.

Deardorff, M.A., Wilde, J.J., Albrecht, M., Dickinson, E., Tennstedt, S., Braunholz, D., Mönnich, M., Yan, Y., Xu, W., Gil-Rodríguez, M.C., et al. (2012). RAD21 mutations cause a human cohesinopathy. Am J Hum Genet 90, 1014–1027.

Dekker, J., and Mirny, L. (2016). The 3D Genome as Moderator of Chromosomal Communication. Cell 164, 1110–1121.

DeScipio, C., Kaur, M., Yaeger, D., Innis, J.W., Spinner, N.B., Jackson, L.G., and Krantz, I.D. (2005). Chromosome rearrangements in cornelia de Lange syndrome (CdLS): report of a der(3 t(3;12)(p25.3;p13.3) in two half sibs with features of CdLS and review of reported CdLS cases with chromosome rearrangements. Am J Med Genet 137, 276–282.

Dheur, S., Saupe, S.J., Genier, S., Vazquez, S., and Javerzat, J.P. (2011). Role for cohesin in the formation of a heterochromatic domain at fission yeast subtelomeres. Mol Cell Biol 31, 1088–1097.

Dorsett, D. (2011). Cohesin: genomic insights into controlling gene transcription and development. Curr Opin Genet Dev 21, 199–206.

Fischer, T., Cui, B., Dhakshnamoorthy, J., Zhou, M., Rubin, C., Zofall, M., Veenstra, T.D., and Grewal, S.I. (2009). Diverse roles of HP1 proteins in heterochromatin assembly and functions in fission yeast. Proc Natl Acad Sci 106, 8998–9003.

Gillespie, P.J., and Hirano, T. (2004). Scc2 couples replication licensing to sister chromatid cohesion in Xenopus egg extracts. Curr Biol 14, 1598–1603.

Gregson, H.C., Schmiesing, J.A., Kim, J.S., Kobayashi, T., Zhou, S., and Yokomori, K. (2001). A potential role for human cohesin in mitotic spindle aster assembly. The Journal of biological chemistry 276, 47575–47582.

Grummt, I. (2013). The nucleolus—guardian of cellular homeostasis and genome integrity. Chromosoma 122, 487–497.

Hammoud, L., Burger, D.E., Lu, X., and Feng, Q. (2009). Tissue inhibitor of metalloproteinase-3 inhibits neonatal mouse cardiomyocyte proliferation via EGFR/JNK/SP-1 signaling. Am J Physiol Cell Physiol 296, C735–745.

Hannan, K.M., Sanij, E., Rothblum, L.I., Hannan, R.D., and Pearson, R.B. (2012). Dysregulation of RNA polymerase I transcription during disease. Biochim Biophys Acta *doi:pii: S1874-9399(12)00185-X*., Epub.

Harris, B., Bose, T., Lee, K.K., Wang, F., Lu, S., Ross, R.T., Zhang, Y., French, S.L., Beyer, A.L., Slaughter, B.D., et al. (2014). Cohesion promotes nucleolar structure and function. Mol Biol Cell 25, 337–346.

Heale, J.T., Ball, J. A. R., Schmiesing, J.A., Kim, J.S., Kong, X., Zhou, S., Hudson, D., Earnshaw, W.C., and Yokomori, K. (2006). Condensin I interacts with the PARP-1-XRCC1 complex and functions in DNA single-stranded break repair. Mol Cell 21, 837–848.

Hetman, M., and Pietrzak, M. (2012). Emerging roles of the neuronal nucleolus. Trends Neurosci 35, 305–314.

Hinshaw, S.M., Makrantoni, V., Kerr, A., Marston, A.L., and Harrison, S.C. (2015). Structural evidence for Scc4-dependent localization of cohesin loading. Elife 4, e06057.

Hisaoka, M., Nagata, K., and Okuwaki, M. (2014). Intrinsically disordered regions of nucleophosmin/B23 regulate its RNA binding activity through their inter- and intra-molecular association. Nucleic Acids Res 42, 1180–1195.

Horsfield, J.A., Print, C.G., and Mönnich, M. (2012). Diverse developmental disorders from the one ring: distinct molecular pathways underlie the cohesinopathies. Front Genet 3, 171.

Huang, K., Jia, J., Wu, C., Yao, M., Li, M., Jin, J., Jiang, C., Cai, Y., Pei, D., Pan, G., et al. (2013). Ribosomal RNA gene transcription mediated by the master genome regulator protein CCCTC-binding factor (CTCF) is negatively regulated by the condensin complex. J Biol Chem 288, 26067–26077.

Kadakia, S., Helman, S.N., Badhey, A.K., Saman, M., and Ducic, Y. (2014). Treacher Collins Syndrome: the genetics of a craniofacial disease. Int J Pediatr Otorhinolaryngol 78, 893–898.

Kagey, M.H., Newman, J.J., Bilodeau, S., Zhan, Y., Orlando, D.A., van Berkum, N.L., Ebmeier, C.C., Goossens, J., Rahl, P.B., Levine, S.S., et al. (2010). Mediator and cohesin connect gene expression and chromatin architecture. Nature 467, 430–435.

Kawauchi, S., Calof, A.L., Santos, R., Lopez-Burks, M.E., Young, C.M., Hoang, M.P., Chua, A., Lao, T., Lechner, M.S., Daniel, J.A., et al. (2009). Multiple organ system defects and transcriptional dysregulation in the nipbl mouse, a model of cornelia de lange syndrome. PLoS Genet 5, e1000650.

Kikuchi, S., Borek, D.M., Otwinowski, Z., Tomchick, D.R., and Yu, H. (2016). Crystal structure of the cohesin loader Scc2 and insight into cohesinopathy. Proc Natl Acad Sci USA 113, 12444–12449.

Kong, X., Ball, A.R., Jr., Pham, H.X., Zeng, W., Chen, H.Y., Schmiesing, J.A., Kim, J.S., Berns, M., and Yokomori, K. (2014). Distinct functions of human cohesin-SA1 and cohesin-SA2 in double-strand break repair. Mol Cell Biol 34, 685–698.

Konig, J., Zarnack, K., Rot, G., Curk, T., Kayikci, M., Zupan, B., Turner, D.J., Luscombe, N.M., and Ule, J. (2010). iCLIP reveals the function of hnRNP particles in splicing at individual nucleotide resolution. Nat Struct Mol Biol 17, 909–915.

Konig, J., Zarnack, K., Rot, G., Curk, T., Kayikci, M., Zupan, B., Turner, D.J., Luscombe, N.M., and Ule, J. (2011). iCLIP--transcriptome-wide mapping of protein-RNA interactions with individual nucleotide resolution. J Vis Exp.

Krantz, I.D., McCallum, J., DeScipio, C., Kaur, M., Gillis, L.A., Yaeger, D., Jukofsky, L., Wasserman, N., Bottani, A., Morris, C.A., et al. (2004). Cornelia de Lange syndrome is caused by mutations in NIPBL, the human homolog of Drosophila melanogaster Nipped-B. Nat genet 36, 631–635.

Laloraya, S., Guacci, V., and Koshland, D. (2000). Chromosomal addresses of the cohesin component Mcd1p. J Cell Biol 151, 1047–1056.

Lechner, M.S., Schultz, D.C., Negorev, D., Maul, G.G., and Rauscher, F.J.r. (2005). The mammalian heterochromatin protein 1 binds diverse nuclear proteins through a common motif that targets the chromoshadow domain. Biochem Biophys Res Commun 331, 929–937.

Li, W., Notani, D., Ma, Q., Tanasa, B., Nunez, E., Chen, A.Y., Merkurjev, D., Zhang, J., Ohgi, K., Song, X., et al. (2013). Functional roles of enhancer RNAs for oestrogen-dependent transcriptional activation. Nature, doi:10.1038/nature12210.

Lin, C.I., and Yeh, N.H. (2009). Treacle recruits RNA polymerase I complex to the nucleolus that is independent of UBF. Biochem Biophys Res Commun 386, 396–401.

Liu, J., and Krantz, I.D. (2009). Cornelia de Lange syndrome, cohesin, and beyond. Clin Genet 76, 303–314.

Mannini, L., Cucco, F., Quarantotti, V., Krantz, I.D., and Musio, A. (2013). Mutation spectrum and genotype-phenotype correlation in Cornelia de Lange syndrome. Hum Mutat 34, 1589–1596.

Martens, J.H., O’Sullivan, R.J., Braunschweig, U., Opravil, S., Radolf, M., Steinlein, P., and Jenuwein, T. (2005). The profile of repeat-associated histone lysine methylation states in the mouse epigenome. The EMBO journal 24, 800–812.

Merkenschlager, M., and Nora, E.P. (2016). CTCF and Cohesin in Genome Folding and Transcriptional Gene Regulation. Annu Rev Genomics Hum Genet 17, 17–43.

Musio, A., Selicorni, A., Focarelli, M.L., Gervasini, C., Milani, D., Russo, S., Vezzoni, P., and Larizza, L. (2006). X-linked Cornelia de Lange syndrome owing to SMC1L1 mutations. Nat Genet 38, 528–530.

Muto, A., Calof, A.L., Lander, A.D., and Schilling, T.F. (2011). Multifactorial origins of heart and gut defects in nipbl-deficient zebrafish, a model of Cornelia de Lange Syndrome. PLoS Biol 9, e1001181.

Narla, A., and Ebert, B.L. (2010). Ribosomopathies: human disorders of ribosome dysfunction. Blood 115, 3196–3205.

Newkirk, D.A., Chen, Y.Y., Chien, R., Zeng, W., Biesinger, J., Flowers, E., Kawauchi, S., Santos, R., Calof, A.L., Lander, A.D., et al. (2017). The effect of Nipped-B-like (Nipbl) haploinsufficiency on genome-wide cohesin binding and target gene expression: modeling Cornelia de Lange syndrome. Clin Epigenetics 9, 89.

Nonaka, N., Kitajima, T., Yokobayashi, S., Xiao, G., Yamamoto, M., Grewal, S.S., and Watanabe, Y. (2002). Recruitment of cohesin to heterochromatic regions by Swi6/HP1 in fission yeast. Nat Cell Biol 4, 89–93.

Oka, Y., Suzuki, K., Yamauchi, M., Mitsutake, N., and Yamashita, S. (2011). Recruitment of the cohesin loading factor NIPBL to DNA double-strand breaks depends on MDC1, RNF168 and HP1γ in human cells. Biochem Biophys Res Commun 411, 762–767.

Parelho, V., Hadjur, S., Spivakov, M., Leleu, M., Sauer, S., Gregson, H.C., Jarmuz, A., Canzonetta, C., Webster, Z., Nesterova, T., et al. (2008). Cohesins functionally associate with CTCF on mammalian chromosome arms. Cell 132, 422–433.

Parlato, R., and Kreiner, G. (2012). Nucleolar activity in neurodegenerative diseases: a missing piece of the puzzle? J Mol Med (Berl) Epub.

Pederson, T. (2011). The nucleolus. Cold Spring Harb Perspect Biol 3, doi:pii: a000638.

Poortinga, G., Wall, M., Sanij, E., Siwicki, K., Ellul, J., Brown, D., Holloway, T.P., Hannan, R.D., and McArthur, G.A. (2011). c-MYC coordinately regulates ribosomal gene chromatin remodeling and Pol I availability during granulocyte differentiation. Nucleic acids research 39, 3267–3281.

Remeseiro, S., Cuadrado, A., Kawauchi, S., Calof, A.L., Lander, A.D., and Losada, A. (2013). Reduction of Nipbl impairs cohesin loading locally and affects transcription but not cohesion-dependent functions in a mouse model of Cornelia de Lange Syndrome. Biochim Biophys Acta 1832, 2097–2102.

Renaud, J.B., Boix, C., Charpentier, M., De Cian, A., Cochennec, J., Duvernois-Berthet, E., Perrouault, L., Tesson, L., Edouard, J., Thinard, R., et al. (2016). Improved Genome Editing Efficiency and Flexibility Using Modified Oligonucleotides with TALEN and CRISPR-Cas9 Nucleases. Cell Rep 14, 2263–2272.

Sakai, D., and Trainor, P.A. (2016). Face off against ROS: Tcof1/Treacle safeguards neuroepithelial cells and progenitor neural crest cells from oxidative stress during craniofacial development. Dev Growth Differ 58, 577–585.

Serrano, A., Rodríguez-Corsino, M., and Losada, A. (2009). Heterochromatin protein 1 (HP1) proteins do not drive pericentromeric cohesin enrichment in human cells. PLoS One 4, e5118.

Shav-Tal, Y., Blechman, J., Darzacq, X., Montagna, C., Dye, B.T., Patton, J.G., Singer, R.H., and Zipori, D. (2005). Dynamic sorting of nuclear components into distinct nucleolar caps during transcriptional inhibition. Molecular biology of the cell 16, 2395–2413.

Takahashi, T.S., Basu, A., Bermudez, V., Hurwitz, J., and Walter, J.C. (2008). Cdc7-Drf1 kinase links chromosome cohesion to the initiation of DNA replication in Xenopus egg extracts. Genes Dev 22, 1894–1905.

Takahashi, T.S., Yiu, P., Chou, M.F., Gygi, S., and Walter, J.C. (2004). Recruitment of Xenopus Scc2 and cohesin to chromatin requires the pre-replication complex. Nat Cell Biol 6, 991–996.

Teng, T., Thomas, G., and Mercer, C.A. (2013). Growth control and ribosomopathies. Curr Opin Genet Dev 23, 63–71.

Tonkin, E.T., Wang, T.J., Lisgo, S., Bamshad, M.J., and Strachan, T. (2004). NIPBL, encoding a homolog of fungal Scc2-type sister chromatid cohesion proteins and fly Nipped-B, is mutated in Cornelia de Lange syndrome. Nat Genet 36, 636–641.

Valdez, B.C., Henning, D., So, R.B., Dixon, J., and Dixon, M.J. (2004). The Treacher Collins syndrome (TCOF1) gene product is involved in ribosomal DNA gene transcription by interacting with upstream binding factor. Proc Natl Acad Sci USA 101, 10709–10714.

van den Berg, D.L.C., Azzarelli, R., Oishi, K., Martynoga, B., Urbán, N., Dekkers, D.H.W., Demmers, J.A., and Guillemot, F. (2017). Nipbl Interacts with Zfp609 and the Integrator Complex to Regulate Cortical Neuron Migration. Neuron 93, 348–361.

Van Nostrand, E.L., Pratt, G.A., Shishkin, A.A., Gelboin-Burkhart, C., Fang, M.Y., Sundararaman, B., Blue, S.M., Nguyen, T.B., Surka, C., Elkins, K., et al. (2016). Robust transcriptome-wide discovery of RNA-binding protein binding sites with enhanced CLIP (eCLIP). Nat Methods 13, 508–514.

Wang, L., and Brown, S.J. (2006). BindN: a web-based tool for efficient prediction of DNA and RNA binding sites in amino acid sequences. Nuc Acids Res 34, W243–248.

Watrin, E., Schleiffer, A., Tanaka, K., Eisenhaber, F., Nasmyth, K., and Peters, J.M. (2006). Human Scc4 is required for cohesin binding to chromatin, sister-chromatid cohesion, and mitotic progression. Current biology: CB 16, 863–874.

Yao, C., Biesinger, J., Wan, J., Weng, L., Xing, Y., Xie, X., and Shi, Y. (2012). Transcriptome-wide analyses of CstF64-RNA interactions in global regulation of mRNA alternative polyadenylation. Proceedings of the National Academy of Sciences of the United States of America 109, 18773–18778.

Yelick, P.C., and Trainor, P.A. (2015). Ribosomopathies: Global process, tissue specific defects. Rare Dis 3, e1025185.

Yokomori, K., and Shirahige, K., eds. (2017). Cohesin and Condensin - Methods and Protocols (Springer).

Yuen, K.C., Xu, B., Krantz, I.D., and Gerton, J.L. (2016). NIPBL Controls RNA Biogenesis to Prevent Activation of the Stress Kinase PKR. Cell Rep 14, 93–102.

Zakari, M., Trimble Ross, R., Peak, A., Blanchette, M., Seidel, C., and Gerton, J.L. (2015). The SMC Loader Scc2 Promotes ncRNA Biogenesis and Translational Fidelity. PLoS Genet 11, e1005308.

Zeng, W., de Greef, J.C., Chen, Y.-Y., Chien, R., Kong, X., Gregson, H.C., Winokur, S.T., Pyle, A., Robertson, K.D., Schmiesing, J.A., et al. (2009a). Specific loss of histone H3 lysine 9 trimethylation and HP1γ/cohesin binding at D4Z4 repeats is associated with facioscapulohumeral dystrophy (FSHD). PLoS Genet 5, e1000559.

Zeng, W., de Greef, J.C., Chen, Y.Y., Chien, R., Kong, X., Gregson, H.C., Winokur, S.T., Pyle, A., Robertson, K.D., Schmiesing, J.A., et al. (2009b). Specific loss of histone H3 lysine 9 trimethylation and HP1gamma/cohesin binding at D4Z4 repeats is associated with facioscapulohumeral dystrophy (FSHD). PLoS Genet 5, e1000559.

Zhang, Y., Sikes, M.L., Beyer, A.L., and Schneider, D.A. (2009). The Paf1 complex is required for efficient transcription elongation by RNA polymerase I. Proc Natl Acad Sci 106, 2153–2158.

Zheng, G., Kanchwala, M., Xing, C., and Yu, H. (2018). MCM2-7-dependent cohesin loading during S phase promotes sister-chromatid cohesion. Elife 7.

Zuin, J., Franke, V., van Ijcken, W.F., van der Sloot, A., Krantz, I.D., van der Reijden, M.I., Nakato, R., Lenhard, B., and Wendt, K.S. (2014). A cohesin-independent role for NIPBL at promoters provides insights in CdLS. PLoS Genet 10, e1004153.

